# Edaphic specialization onto bare, rocky outcrops as a factor in the evolution of desert angiosperms

**DOI:** 10.1101/2022.09.27.509613

**Authors:** Isaac H. Lichter Marck, Bruce G. Baldwin

**Author notes:** Isaac H. Lichter Marck, **Email:**. **Author Contributions:** ILM & BGB designed the study; ILM carried out the study; ILM wrote the manuscript. All authors revised and reviewed the manuscript.

## Abstract

Understanding the processes that enable organisms to shift into more arid environments as they emerge is critical for gauging resilience to climate change, yet these forces remain poorly known. In a comprehensive clade-based study, we investigate recent shifts into North American deserts in the rock daisies (Perityleae), a diverse tribe of desert sunflowers (Compositae). We sample Perityleae across two separate contact zones between tropical deciduous forest and desert biomes in western North America and infer a time-calibrated phylogeny based on target capture sequence data. We reconstruct biome shifts using Bayesian inference with paleobiome-informed models and find evidence for seven independent shifts into desert habitats since the onset of aridification in the late Miocene epoch. The earliest shift occurred out of tropical deciduous forests and led to an extensive radiation throughout North American deserts that accounts for the majority of extant desert Perityleae. Reconstructions of life history and micro-habitat in Perityleae reveal a correlation between a suffrutescent perennial life history and edaphic endemism onto rocky outcrops, an ecological specialization that evolved prior to establishment and diversification in deserts. That the insular radiation of desert rock daisies stemmed from ancestors pre-adapted for dry conditions as edaphic endemics in otherwise densely vegetated tropical deciduous forests in northwest Mexico underscores the crucial role of exaptation and dispersal for shifts into arid environments.

**Significance Statement:** The environmentally stressful conditions found in desert regions have often been implicated as the main factor in the evolution of drought tolerance in desert plants. Yet many iconic desert plant lineages evolved prior to the recent emergence of widespread arid climates, suggesting an important role for pre-adaptation (exaptation). In the desert rock daisies (Perityleae), we provide empirical support for this view by showing that life history evolution associated with their ecological specialization onto rock outcrops was a precursor to their establishment and extensive diversification in North American deserts. We caution against assuming the presence of ancient dry biomes based on time-calibrated phylogenies and we emphasize the fundamental roles that exaptation and dispersal play during community assembly in novel environments.

## Introduction

Deserts are the most widespread biomes on earth and predicted to grow in extent as a result of climate change, but the potential for biotic communities to adapt to new levels of increased aridity is poorly understood (1–2). Compared to other biomes, many deserts emerged recently on a geological timescale (3); therefore, their endemic biota provides an ideal system for understanding the organismal potential to fill novel environments created by future climate change. North American (NA) deserts cover most of the southwest U.S. and northern Mexico (4) and appeared for the first time in scattered, ephemeral pockets as recently as the late-Miocene during Neogene uplift (5-7 Ma) and then expanded during the Pliocene (2.4-5 Ma) (5). Despite their recent origin, NA deserts contain a diverse and endemic flora and fauna endowed with striking abilities to escape or tolerate their harsh, arid climate whose origins have been a topic of debate among evolutionary biologists for decades (6–7). Examples of evolutionary adaptation showcased by desert plants include the ephemeral annual lifestyle, and the many types of unique, stunted, thorny, or succulent vegetative growth forms that are features well-suited for drought tolerance or escape (4,8). Most commonly, these features are attributed to rapid adaptive evolution in response to the environmentally stressful conditions that characterize deserts: unpredictable precipitation, extremes of temperature, and coarse, rocky soils (9). Yet this explanation has been difficult to reconcile with the recent emergence of deserts and the age of origin of iconic desert plant lineages in the Oligocene supported by time-calibrated phylogenies and the fossil record (10–12). An alternative explanation is that desert plants evolved by exaptation in arid micro-sites, such as rain-shadows or rock outcrops, within the moist forests of older biomes, and then spread and diversified when the climate became cooler and drier (7). Thus, transitions into desert biomes may have been accomplished by rapid adaptive evolution, or organisms pre-adapted to novel conditions within neighboring biomes may have dispersed into arid environments as they emerged (13).

While the phylogenetic and statistical tools needed to tease apart the roles of adaptive evolution and dispersal during biome shifts have grown substantially in recent years, testing these alternative hypotheses continues to pose a challenge because of the difficulty in pinpointing when and where evolutionary events of particular importance happened from contemporary data (14). Phylogenetic studies set within a paleo-biome and paleo-geographically informed framework have the potential to resolve when and where biome shifts have occurred in the past (15). Yet rapid adaptation or exaptation, by definition, imply both an evolutionary transition in form as well as a shift in ecological context. Only a comprehensive clade-based study integrating both extrinsic (environmental) and intrinsic (trait) data into a time integrated context can provide the varied lines of evidence needed to evaluate hypotheses about the mechanisms underlying biome shifts (14). Here, we investigate biome shifts into NA deserts using a uniquely comprehensive study of the rock-daisy tribe (Perityleae) of the sunflower family (Compositae). We reconstruct shifts into desert biomes using Bayesian inference with paleo-biome informed models, then draw upon unique life history traits closely tied to microclimate specificity to build an integrative understanding of ecological transitions and trait evolution through time for 85 species of Perityleae and close relatives.

The evolution of the megadiverse sunflower family is closely tied to the emergence of arid biomes worldwide, but compared to other dryland plant groups, such as legumes, succulents, or grasses, few studies have examined ecological transitions to arid climates in desert Compositae (10,16–18). Perityleae are one of the most diverse groups of Composites found in the NA deserts and their range extends to tropical deciduous forests adjacent to desert in northwest mainland Mexico and the Baja California peninsula (19–20). Disjunct populations are also present in the Atacama Desert in Chile (19–20). Within the NA deserts, Perityleae show characteristics suggestive of an insular radiation, including high species diversity, geographic speciation, divergent life history strategies, and specialization into disparate ecological niches (20–21). A remarkable aspect of this group’s natural history is the extreme edaphic specialization demonstrated by some taxa, which grow exclusively on bare, rocky outcrops, including steep canyon walls and cliffs on the slopes of desert mountains. Associations with bare rock are exhibited in both desert and tropical habitats, where they appear to be facilitated by a specialized life history trait, a perennating woody caudex lodged into cracks or crevices from which herbaceous shoots resprout annually (suffrutescent perennial life history, Fig. 1). Other members of Perityleae lack this trait and these may be multi-branched woody shrubs, herbaceous perennials, or annuals, including some of the most abundant ephemeral wildflowers (e.g. *Perityle emoryi* Torr., *Perityle californica* Benth., *Galinsogeopsis spilanthoides* Sch. Bip.) found in the seasonal blooms of low deserts in favorable years (19–20).

**Figure 1.**
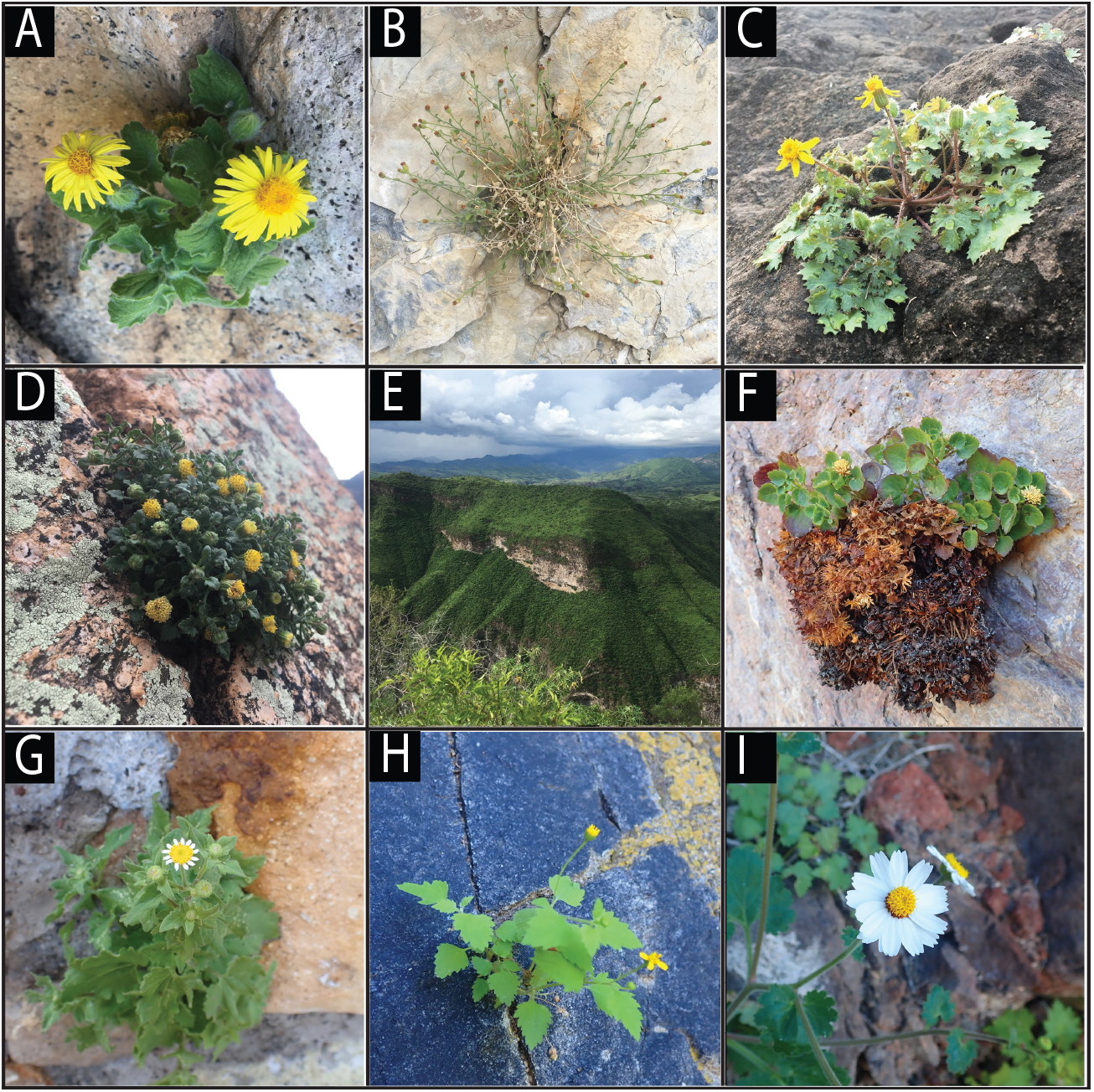
Habit and habitat diversity of Perityleae. (A) *Laphamia cordata*; (B) *Laphamia villosa*; (C) *Laphamia lobata*; (D) *Laphamia cochisensis*; (E) tropical deciduous forest with bare, rocky outcrops in the Sierra Madre Occidental in Sonora, Mexico; (F) *Laphamia cernua*; (G) *Galinsogeopsis spilanthoides*; (H) *Perityle cuneata* var. *cuneata*; (I) *Perityle rotundifolia*; Photos by ILM.

The ecogeographic range, breadth of ecological specializations, and life history variation exhibited by Perityleae makes them an ideal group to study the roles of adaptive evolution and dispersal on biome shifts into the NA deserts; however, a densely sampled phylogeny of this lineage has only recently become available (21). A robust phylogeny for Perityleae based on target capture using the Conserved Compositae Ortholog set (COS) (22) formed the basis for recent re-classification of the group into 9 genera (19). Phylogenies based on target capture data have proved useful in resolving relationships in this group at a variety of scales, but more sampling is needed to fill gaps in previous phylogenies. Here, we describe the results of the most densely sampled phylogenetic study of Perityleae to date based on target capture of >95% of recognized taxa and employ our improved resolution of relationships to test hypotheses regarding shifts into desert biomes. Specifically, we test whether evolutionary shifts to key functional traits for enduring arid conditions occurred as an adaptive response to the environmentally stressful barriers presented by desert habitats, or whether Perityleae pre-adapted (exapted) for arid conditions on arid microsites within older biomes dispersed into deserts with corresponding trait evolution already in place.

## Results and Discussion

### Novel insights into the evolutionary and biogeographic history of Perityleae

Phylogenies of Perityleae based on a concatenated data matrix of 1,419,391 bp of orthologous nuclear loci were highly resolved at most nodes for both maximum likelihood (ML) and Bayesian inference (BI) approaches and congruent with each other, as well as with previously published phylogenies of Perityleae based on COS loci *(SI Appendix,* Figs. S2, S5-6; 55). Sampling of almost all recognized taxa allowed for a comprehensive understanding of evolutionary relationships among Perityleae (*SI Appendix*, Fig. S2). Conspecific samples clustered together for most taxa, but many were found to be non-monophyletic, suggesting the presence of multiple cryptic lineages in need of taxonomic follow-up. Lack of resolution within individual loci or discordance due to ILS and introgressive hybridization was in evidence in our pseudocoalescent phylogenetic analysis, with a major topological difference between the pseudocoalescent tree and the concatenated analysis being the placement of the widespread allopolyploid *P. emoryi*, which resolves as non-monophyletic and sister (in part) to *Laphamia* in our pseudocoalescent tree but as nested in *Perityle* with high support in all other analyses (*SI Appendix*, Fig. S7).

In our reduced sample, maximum clade credibility time-calibrated phylogeny, the root node corresponding to the most recent common ancestor of Perityleae+Eupatorieae and *Helianthus annuus* L. was inferred to have a median age of 15.8033 Ma [9.6371– 20.8277 Ma, 95% HPD] and agrees with the range of times found in previous dating analysis of Heliantheae s.l. (23–24, *SI Appendix*, Fig. S3). A basal split between the two major lineages of Perityleae is inferred at 12.5643 Ma [7.7409–17.1751 Ma, 95 % HPD] and shows remarkable correspondence in timing with estimates of splitting of the Baja California peninsula from mainland Mexico (25). Indeed, a cladogenetic vicariance event resolved at this node separates the principal radiations of Perityleae, which have distributions centered on mainland Mexico and Baja California, respectively (*SI Appendix*, Fig. S4). Recent paleo-biome reconstructions based on fossil, palynological, and imputed climatological data infer a range of first emergence of widespread desert habitats in western NA ~5–7 ma, with climatic cooling and aridification leading to further spread of arid environments during the Pliocene (5). For the most part, divergence events separating the major clades of Perityleae predate widespread climatic aridification and occur prior to dispersal into the Basin and Range Province within more densely vegetated areas of northwest Mexico (Figs. 2, 3). This includes *Laphamia,* which diverged from *Galinsogeopsis* 8.7917 Ma [5.3261–12.0393 Ma, 95% HPD] yet includes a nested, well-supported clade that contains most desert Perityleae, and whose MRCA corresponds with the widespread emergence of deserts at 4.9966 Ma [3.0599– 6.9604 Ma, 95% HPD] (Fig. 2, *SI Appendix*, Fig. S3).

**Figure 2.**
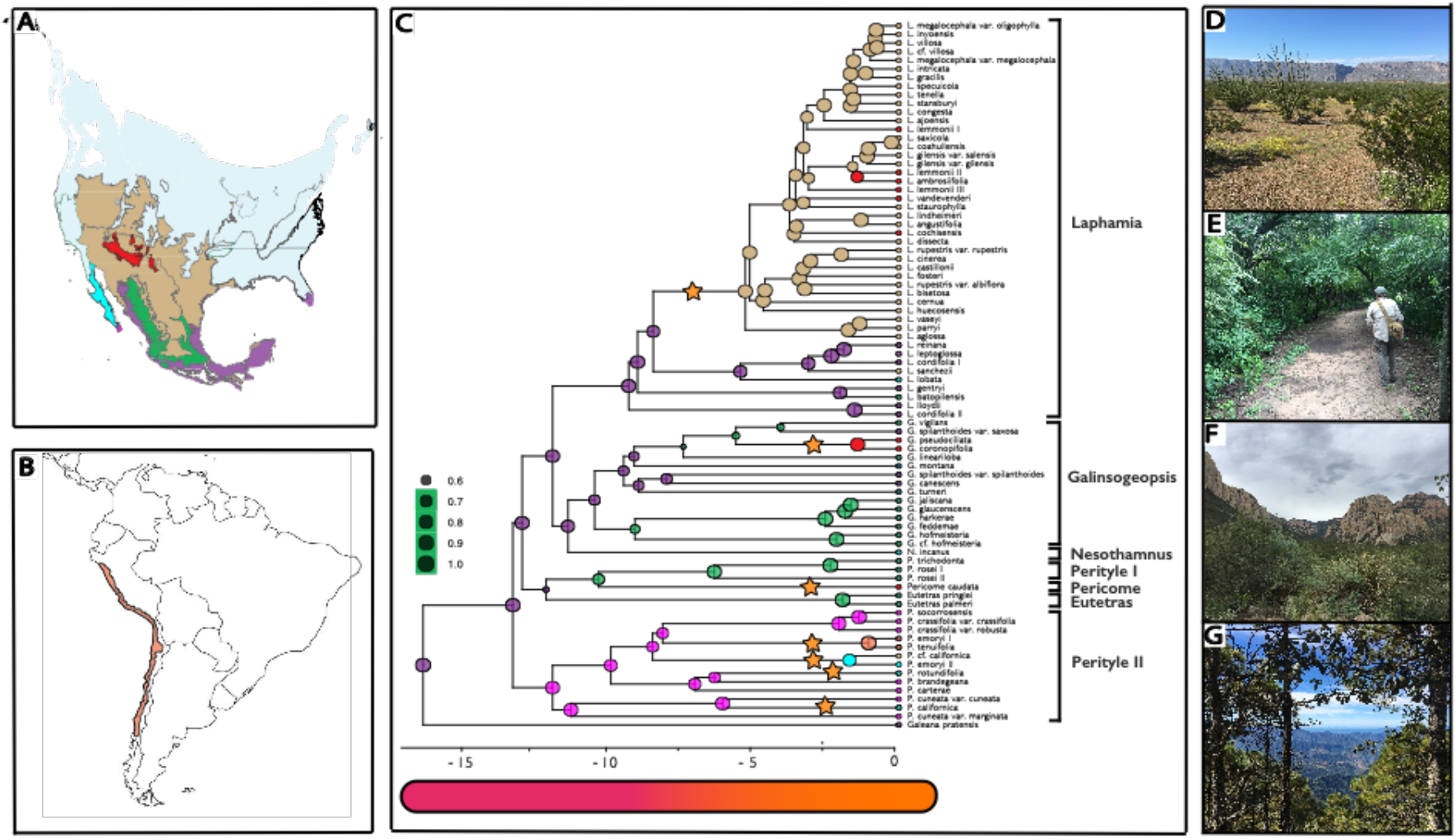
A) Biome map of North America illustrating the biomes addressed in this study, including desert in the Basin and Range Province (brown) and Baja California (blue), subtropical forest in the Basin and Range Province (red) and Trans-Mexican Volcanic Belt of Mexico (green), and tropical deciduous forest in Baja California (pink) and northwest Mexico (purple). (B) Distribution of the geographically disjunct Atacama Desert (orange) in South America. (C) Maximum a posteriori reconstruction of ancestral biome and geographic region in Perityleae inferred using the biome-shift model of Landis et al. (8). Colors at nodes indicate biome-region combinations corresponding to colors in (A) and (B) and the size of the circle its posterior probability. Shifts into the desert biome are indicated with a star. (D-G) Landscape photographs of dominant vegetation types in biomes included in this study. (D) Desert plain in Big Bend National Park in Texas, U.S.A. (E) Tropical deciduous forest in the Sierra de Alamos in Sonora, Mexico. (F) Subtropical woodland in the Chiricahua Mountains of Arizona, U.S.A. (G) Temperate Forest in the Sierra Madre Occidental of Durango, Mexico.

**Figure 3.**
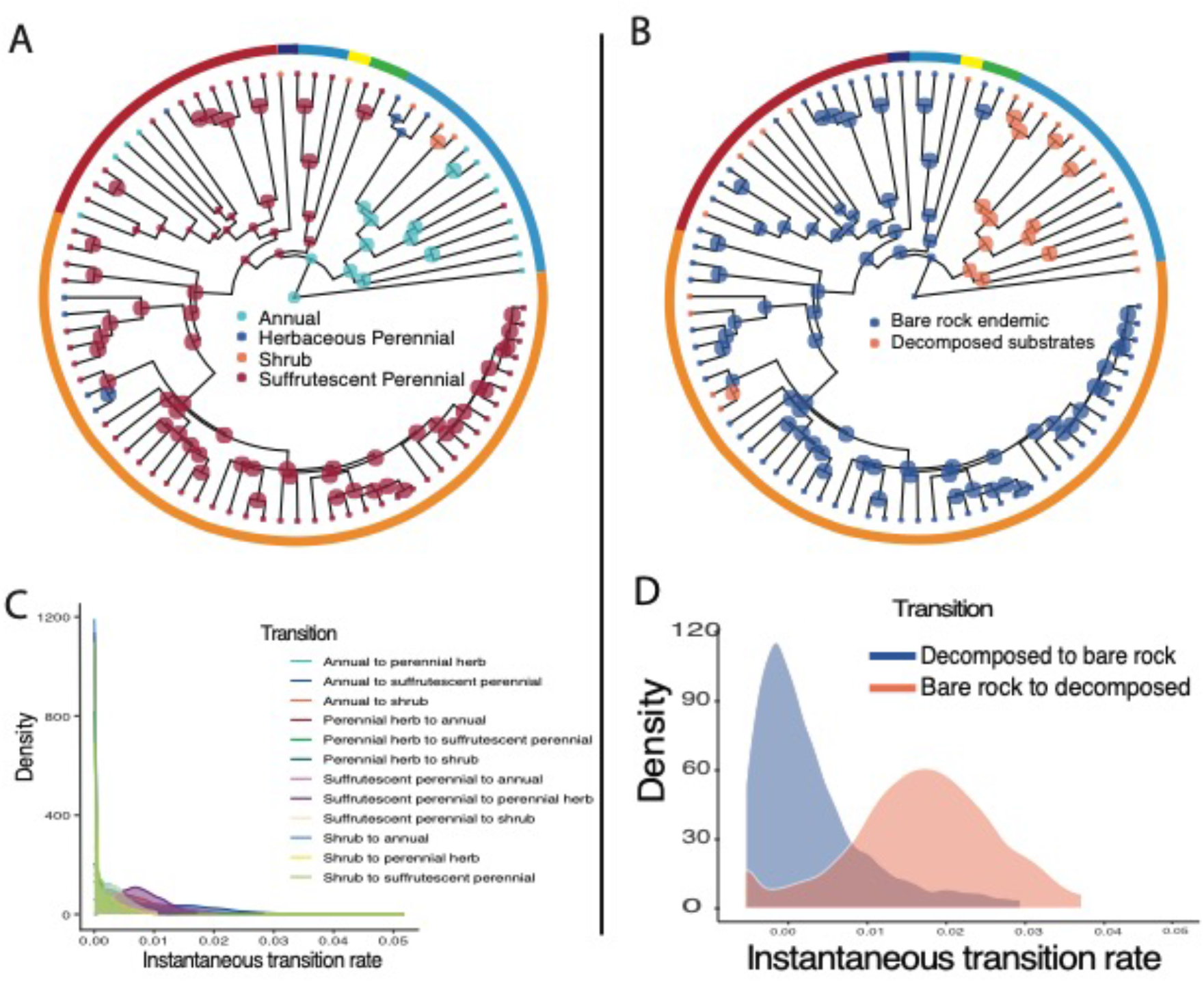
Evolution of edaphic specialization onto bare, rocky outcrops is closely correlated with a suffrutescent life history and evolved early in the evolutionary history of Perityleae. (A) Model-averaged maximum a posteriori (MAP) ancestral state reconstruction of the woody shrub, suffrutescent perennial, herbaceous perennial, and annual life history based on 28,500 iterations of post-burn-in rjMCMC in RevBayes; (B) MAP ancestral state reconstruction of strict edaphic endemism onto bare rock in Perityleae based on 28,500 iterations of post- burn-in rjMCMC in RevBayes; (C) posterior density estimates of transition rate asymmetries between life histories (D) posterior density estimates of transition rate asymmetries between obligate growth on bare rock and occurrence on decomposed substrates.

### Evolution of edaphic specialization onto bare rocky outcrops and life history

Using reversible jump (rj) MCMC with stochastic character mapping to reconstruct the timing and tempo of life history evolution showed rate asymmetries, with transitions from suffrutescent perennials to shrubs, perennial herbs, or annuals inferred as moderate, but transitions to the suffrutescent perennial life history inferred as not significantly different from zero, suggesting irreversibility (Table 1). Model-averaged reconstructions of ancestral states on the maximum a posteriori (MAP) chronogram clarify that the suffrutescent life history evolved once early in the evolutionary history of Perityleae and is a shared ancestral characteristic of the clade containing *Eutetras, Laphamia, Galinsogeopsis*, and *Pericome*, although multiple independent transitions from suffrutescent perennials to shrubs, herbaceous perennials, and annuals occurred more recently within this group (Fig. 3). The MRCA of the mostly Baja California *Perityle* clade is resolved as herbaceous, with a woody shrub life history secondarily derived coincident with dispersal to island or island-like habitats in three cases: in *Perityle carterae, Perityle socorrensis*, and *Perityle tenuifolius*.

**Table 1.** Model parameters derived from ancestral state reconstructions of life history and edaphic affinity using rjMCMC in RevBayes. Bayes Factors are raw proportions of iterations in favor of a non-zero transition rate.

| Character | Transition | Mean transition rate | 95% HPD interval | Bayes Factor |
| --- | --- | --- | --- | --- |
| Life history | Shrub to suffrutescent | 1.3925E-3 | [0,6.467E-3] | 0.509 |
|  | Shrub to herbaceous | 1.3376E-3 | [0,0.0277] | 0.503 |
|  | Suffrutescent to shrub | 3.8089E-3 | [0,0.0204] | 0.904 |
|  | Suffrutescent to herb | 0.0185 | [7.0696E-3,0.03] | 1 |
|  | Herb to suffrutescent | 4.6127E-3 | [0,0.0119] | 0.947 |
|  | Herb to shrub | 7.1473E-3 | [0,0.322] | 1 |
| Edaphic affinity | Bare rock endemic to decomposed substrates | 4.3876E-3 | [0,0.035] | 1 |
|  | Decomposed to bare rock endemic | 0.0191 | [8.6597E-3,0.0312] | 0.947 |

Mirroring patterns in life history evolution, edaphic endemism onto bare rocky outcrops is resolved as the historical ecology of the MRCA of the clade containing *Eutetras, Laphamia, Galinsogeopsis*, and *Pericome*, whereas growth on decomposed substrates is resolved as the ancestral condition for genus *Perityle* (Fig. 2). RjMCMC inferred transition rates were not significantly different from zero for transitions onto bare rocky substrates, suggesting that edaphic endemism may have been lost many times during the evolution of Perityleae, but once lost, was not readily regained (Fig. 3, Table 1). The timing and tempo of transitions from a suffrutescent life history were mirrored by the evolution of edaphic specialization across Perityleae, hinting at a strong evolutionary dependence between obligate growth on bare rock and the perennating woody caudex that characterizes suffrutescent perennials. Bayesian model testing of dependent [Log-likelihood = −51.090204] and independent [log-likelihood = −65.153536] models of evolution of edaphic endemism and a perennating woody caudex supported this relationship [log Bayes Factor = 28.12226], suggesting very strong empirical evidence for evolutionary dependence between these characters across the phylogeny.

### Ecological shifts inferred with paleo-biome informed models

Transition rates between desert, tropical, subtropical, and temperate biomes show asymmetries across our study group with the highest transition rates observed from out of tropical and into desert and subtropical biomes, and the lowest a shift from tropical to temperate biomes (Table 2). Perityleae underwent six independent shifts into desert biomes from two geographically separate source areas (Fig. 2). Three of these occurred from out of the Baja California Peninsula and three from the Trans-Mexican Volcanic Belt, Sierra Madre Occidental, and Pacific plain of northwest Mexico (Fig. 3). Shifts to desert occurred from tropical habitats in at least four cases and a subtropical biome in at least one case. Disjunct populations of Perityleae in the Atacama Desert in South America arrived via long-distance dispersal that coincided with a biome shift from tropical environments in southern Baja California and the neighboring Revillagigedo archipelago. The ecological origin of desert dwelling *Galinsogeopsis ciliata* and *G. coronopifolia* was equivocal [subtropical origin PP = 0.5452, tropical origin PP = 0.4122].

**Table 2.** Results of a biome-shift analysis quantifying relative dependence on three separate underlying q matrices: uninformed (w_u), paleo-geographically informed (w_g), and paleo-biome informed (w_b), as well as non-paleo-biome informed (w_not_b).

| Statistic | Relevance | Mean | 95% HPD interval |
| --- | --- | --- | --- |
| w_u | Transition matrix uninformed by paleo-biome or paleo-geography | 0.303 | [0.0146,0.6481] |
| w_g | Relative importance of paleo-geography | 0.2471 | [4.3735E-4,0.5846] |
| w_b | Relative dependence on paleo-biome<br>structure | 0.4496 | [0.0419,0.8315] |
| w_not_b | Independence<br>from paleo-biome structure | 0.5504 | [0.1685,0.9581] |

The vast majority (~57 spp, >85%) of desert dwelling Perityleae stem from a single shift into the NA deserts from tropical deciduous forests in northwest Mexico shortly after the onset of aridification in the late Miocene (Fig. 2). In contrast, remaining shifts into a desert climate are resolved as recent, occurring in terminal branches with little contemporary lineage diversification. Estimation of transition rates between biomes within a time-integrated model of biome availability and adjacency suggested only moderate dependence of shifts on increasing availability of desert habitats following late Miocene aridification [w_b = 0.45, w_u = 0.303, and w_g = 0.247] (Table 2). Bayesian model testing using rjMCMC to compare dependence on increasing biome availability rejected dependence on paleobiome structure [log Bayes Factor = −10], suggesting support for a model in which shifts into dry habitats were independent from the increase in aridity through time.

### Edaphic specialization onto bare rocky outcrops as a precursor to establishment and diversification in a desert biome

The variable and extreme conditions found in deserts present great challenges for plants, with the lack of water and coarse, rocky soils generally the most challenging conditions plants face (6–9). These environmentally stressful conditions in desert regions have long been regarded as the stimulus for adaptive strategies for drought tolerance (6,26). Yet the time-calibrated phylogenetic studies of many desert plant lineages that have been carried out to-date conflict with this conventional view, showing instead that major floristic elements of arid ecosystems worldwide, including cacti, legumes, ocotillos, elephant trees, and others, acquired their unique morphologies prior to the emergence of widespread dry climates in the Neogene and then radiated synchronously during recent onset of aridification (10, 27, SI Appendix). This pattern has caused some to speculate that arid biomes appeared much earlier than previously believed, as early as in the Eocene (~20 million years before the mid-Miocene) (17, 28). How else could these traits have evolved without the harsh, arid conditions that produced them? We provide support for an alternative hypothesis, that since many plants living in a desert environment today did not originally evolve in a desert setting, features seen in each plant may not be adaptations to desert living but carryovers from some other set of conditions. Specifically, we find that rock daisies of genus *Laphamia* evolved first as edaphic endemics on “micro deserts” of bare rocky outcrops within otherwise densely vegetated environments, and then spread and diversified more recently under cooler, drier climatic conditions. Most of the ca. 58 minimum-rank taxa of Perityleae in *Laphamia* that inhabit the NA deserts show conserved life history evolution associated with ancestral ecological specialization. Dispersal into the Basin and Range Province when arid conditions intensified in the mid-Miocene evidently led to the ecological release and rapid radiation of *Laphamia* onto ecological islands, including bare, edaphic islands and decomposed substrates, scattered across the varied landscapes of the NA deserts (*SI Appendix*, Fig. S1).

The idea that traits conferring drought tolerance may have been a prerequisite rather than a consequence of establishment in the NA deserts is not new (7,29), yet several non-mutually exclusive lines of evidence suggest it may be a scenario of broader applicability than previously recognized in explaining the enigmatic origins of desert biota. Within forested landscapes, rocky outcrops tend to form micro-habitats that consistently harbor a more aridity-resistant flora than their surroundings, and often contain disjunct populations of plants from adjacent, or distant, more open, dry biomes (30). Rocky outcrops in the tropical deciduous forests that are adjacent to the NA deserts and considered a major source area for expansions into those deserts, harbor familiar members of diverse desert plant and animal groups, including cacti, agaves, euphorbias, rattlesnakes, and lizards (7, *SI Appendix*, Figure S1). In these and other forested environments, edaphic outcrops have presented a stable, continually present, and widespread nucleus for the evolution of drought tolerance since the Cretaceous era; in contrast, widespread dry habitats have been rare throughout time, especially during the cool, moist periods of the Oligocene and early Miocene epochs (3).

Growth in edaphically arid substrates poses great challenges for plants and many of the adaptive strategies that confer tolerance to stressful soils overlap with those generally considered useful for coping with stress due to aridity (31–32). These traits include perennating organs; a dense indument; reduction in leaf area; deciduousness; aphylly; tolerance of exposure to UV, intense fluctuations in temperature, wind, and lack of facilitating nurse plants; as well as enhanced herbivore defense strategies to compensate for the high cost of regeneration after attack (32–33). Transitions onto edaphically stressful soils have often been implicated in driving rapid and drastic plant evolution in growth form because of the tradeoffs for resource availability and defense that manifest in nutrient poor conditions (34–35). Tolerance for bare soils has also been implicated in the evolution of further specialization onto chemically unusual serpentine soils, suggesting that bare soils may represent an important hurdle that, once crossed, enables expansions into other stressful niches (32).

Edaphic exaptation as a factor in the evolution of desert angiosperms remains to be reconciled with prevalent theories regarding the selective pressures that maintain edaphic endemism as a stable ecological strategy in dry environments. Most edaphic endemics, including those in Perityleae, can grow and thrive on zonal (normal) soils in greenhouse conditions, suggesting that tradeoffs in competitive ability or herbivore defense play a crucial role in excluding them from zonal soils in a natural community context (32). In the dense forests where Perityleae evolved edaphic endemism onto bare rocky outcrops, competitive exclusion provides a plausible explanation for narrow endemism onto more stressful bare rock, but in contemporary deserts, where the vegetation is open, and facilitation by nurse plants common, escape from competition does not provide a satisfactory explanation for edaphic endemism, suggesting constraints associated with specialization. Another promising explanation is the inverse texture hypothesis (36), which states that in wet ecosystems, where nutrients are more limiting than water, finer, more fertile soils are more productive than coarse, nutrient-poor substrates. However, in dry ecosystems coarser soils allow water to infiltrate below the evaporative zone more rapidly than finer soil, allowing them to retain soil moisture longer. Tests of the inverse soil texture hypothesis are scarce, but, if true, rocky substrates could constitute a crucial biome bridge, facilitating shifts into more arid adaptive zones without fundamental changes to a group’s conserved ecological niche.

## Conclusion

Our knowledge of biodiversity remains fragmentary, with diverse groups of specialists often being the least well-documented facets of a given ecosystem. Ecological specialists have been considered evolutionary dead-ends (37), but here we find evidence for the successful spread and radiation of a group of edaphic endemics from the tropics into the open, arid habitats of the NA deserts. These data not only highlight the NA desert biome as an important area for in situ plant diversification, but they also add to recent studies (32, 38–39) recognizing the major roles that resource tracking and pre-adaptation (exaptation) play in generating biodiversity throughout space and time. More importantly, this research highlights the importance of dispersal, especially considering the potentially limited responses of plants to climate change towards greater aridity (40). Furthermore, this research underscores the need to document diverse groups of ecological specialists, not only because of their heightened vulnerability to habitat loss and global change, but for their potential to mediate future biotic responses to the emergence of novel and extreme climates.

## Materials and Methods

### Sampling, sequencing, phylogenetic inference, and divergence time estimation (DTE)

We carried out sequence capture using the COS baits (22–23) for target capture of ~1060 low copy nuclear loci for 114 samples encompassing all minimum-rank taxa of subtribe Peritylinae except for seven taxa for which material was unattainable or sequence yield insufficient, plus three outgroup taxa (*SI Appendix*, Table S1). We extracted DNA, enriched libraries, and processed data using the approach of ref. 21, with the exception that we used iterative reference-based assembly in the HybPiper pipeline (41). We used HybPiper’s paralog investigator script to identify and exclude potentially problematic loci and the intronerate.py script to obtain supercontigs, which included exons and their overlapping flanking introns (see 65 for details). Loci with less than 95% occupancy were removed and the remaining 229 loci aligned using MAFFT (42).

ML and BI trees were generated using RAxML-NG (43) and ExaBayes (44) on the CIPRES science supercomputing portal (45) and a pseudocoalescent tree from gene trees while accounting for ILS was produced using ASTRAL-III (46) (*SI Appendix*, Figs. S2, S5-7). A dataset for DTE with one representative of each taxon was generated by choosing the sample that conformed most in terms of geography and morphology to the type, retaining individuals of each independent evolutionary lineage when taxa were found to be non-monophyletic. To this constrained topology, a time-calibrated phylogenetic hypothesis was obtained using Bayesian inference in MCMCtree (47) using a subset of loci and a secondary calibration of the 95% HPD interval estimate of the split of *Helianthus annuus* from Eupatorieae + Perityleae of 7.93 to 23.22 ma, applied to the root of our phylogeny with a uniform prior (24) (*SI Appendix*, Fig. S3).

### Evolution of life history and edaphic specialization

Observations of life history and edaphic specialization came from herbarium label data, taxonomic treatments, species descriptions, and the 1^st^ author’s fieldtrips to the southwest U.S. and northern Mexico during 2017, 2018, and 2019 (19–21). For the purposes of this study, herbaceous plants included those that have an intra-annual or perennial lifecycle and lack woody tissue. Suffrutescent perennials resprout annually from a woody stem base, while shrubs are multi-stemmed and woody above ground. Bare rock surfaces were defined as cliffs, boulders, rocky outcrops, or canyon walls of solid consistency with minimal soil accumulation outside of cracks and crevices. Taxa were considered bare rock endemics if all known occurrences were on solid rock substrates. Taxa that were not endemic to bare rock grew on decomposed substrates, which were defined as various weathered soils that fracture on contact and are composed of gravel, sand, silt, or clay. We estimated rates of transitions and inferred ancestral states for life history and edaphic specialization using Bayesian model testing with rjMCMC and stochastic character mapping in RevBayes (48). To test for associations between life history and endemism on rocky outcrops we used BayesTraits to compare the fit (log-likelihood) of two continuous-time Markov models of life history and edaphic evolution, a dependent and independent model (49).

### Historical biogeography and ecology with paleo-biome informed models

To reconstruct the timing and tempo of biome shifts we used the Biome-Shift model developed by Landis et al. (15) and implemented in RevBayes (48) to consider all combinations of four geographical regions (South America, Baja California, the Trans-Mexican Volcanic Belt, and the Basin and Range Province) with four biomes (tropical, temperate, subtropical, and desert). Ancestral biogeographic ranges were additionally inferred and compared to our paleo-biome informed model using a Bayesian implementation of a DEC model in RevBayes (48,51) (*SI Appendix*, Fig. S4). Our Biome-Shift model was structured with four epochs: Oligocene (33.9-23.0), early Miocene (23.0-16), mid/late Miocene (16-5.3), and recent (5.3-0). South America was coded as having zero connectivity to any other areas in all epochs. We also modeled the split of the Baja California Peninsula from mainland Mexico beginning ~12 Ma (25) by assigning strong connectivity to the Basin and Range Province and Trans-Mexican Volcanic Belt through the Oligocene and early Miocene, weak connectivity in the mid/late Miocene, and zero connectivity in the recent epoch. Sizes and distributions of temperate, tropical, and subtropical biomes were modeled as constant from the Oligocene to present and desert biomes modeled as emerging with weak connectivity in the late Miocene and strong connectivity in the recent epoch.

Additional details of the methods are available in *SI Appendix*.

## Acknowledgments and funding sources

We thank A. Michael Powell, the late Vicki A. Funk, and the late Harold Robinson for their mentorship of ILM, personnel from the following herbaria for specimen access: AZ, MO, NY, US, CIIDIR, USON, CUCBA, UC/JEPS, SRSU, and CAS, and Sophia Winitsky, Jesus E. Sanchez, Pablo Carillo-Reyes, Arturo Castro Castro, Jon Rebman, Robert Villa, Jesus Pablo Carillo, Gabriel Johnson, Bridget Wessa, Lydia Davis, Michael Landis, Michael May, Carrie Tribble, Will Freyman, and Carolina Siniscalchi for assistance. Funding was provided by the Smithsonian Institution Fellowship Program, Philomathia Foundation, Department of Integrative Biology, UC Natural Reserve System, Society of Systematic Biologists, American Society of Plant Taxonomists, California Native Plant Society, California Botanical Society, Southern California Botanists, and the Lawrence R. Heckard Fund of the Jepson Herbarium. The authors declare no conflict of interest.

