## supplemental file for "Edaphic specialization onto bare, rocky outcrops as a factor in the evolution of desert angiosperms"

[*] Isaac H. Lichter-Marck

**This PDF file includes:**

Supplementary text
Figures S1 to S8
Table S1
SI References

**Supplemental Materials and Methods**

**Sampling, sequencing, phylogenetic inference, and divergence time estimation (DTE).**

Our study included near-comprehensive coverage of all minimum-rank taxa of tribe Perityleae except *Pericome macrocephala* B.L. Rob., *Perityle aurea* Rose, *Laphamia grandifolia* (Brandegee) I.H. Lichter-Marck, *L. warnockii* (A.M. Powell) I.H. Lichter-Marck, and the four taxa of genus *Villanova* Lag. in subtribe Galeaninae for which material was unattainable. Three additional taxa, *Laphamia ciliata* Dewey, *L. quinqueflora* Steyermark, and *L. vitreomontana* (Warnock) I.H. Lichter-Marck were excluded from this study due to low sequence yield. As outgroup taxa, we included publicly available sequence data from GenBank sequence read archive (SRA) for *Helianthus annuus* L. of tribe Heliantheae and *Conoclinium coelestinum* DC. and *Eutrochium fistulosum* (Barratt) E.E. Lamont of tribe Eupatorieae. We extracted DNA from silica dried or herbarium leaf tissue using the DNeasy Plant Mini Kit (Qiagen, Valencia, CA, USA) or a modified version of Doyle and Doyle's (1) CTAB protocol identical to the one used by Lichter-Marck et al. (2). We prepped libraries using the HyperPrep protocol (Kapa Biosystems Inc., Wilmington, MA) in quarter reactions. Libraries were pooled 8 per reaction and enriched for 1060 low-copy nuclear genes using the COS kit (3-5) according to the directions provided by myBaits (Arbor Biosciences, Ann Arbor, MI). Enriched libraries were sent to Genewiz (South Plainfield, NJ) for massively parallel paired-end 150 bp sequencing on a HiSeq 4000, yielding >91,000 MB of raw sequencing data. Raw sequencing yields were deposited in the GenBank SRA under bioproject PRJNA###. See Table S1 for voucher and sample SRA information.

Maximum likelihood (ML) and Bayesian inference (BI) trees were generated in RAxML-NG (6) and ExaBayes (7) using a partitioned data matrix, with each locus independently assigned a GTR+Gamma+I model of molecular evolution. The best ML tree was selected after 1000 independent searches and 1000 bootstrap replicates. The BI analysis used two independent chains to sample the posterior distribution of trees over 100,000 generations before computing a maximum clade credibility (MCC) tree based on a 50% majority rule. Gene trees used in the ASTRAL analysis were inferred separately using RAxML-NG with a GTR+Gamma+I substitution model and nodes with less than 20% bootstrap support collapsed (8). Our MCMCtree analysis used a HKY model of molecular substitution with four gamma distributed rate categories and a relaxed molecular clock conditioned on a log normally distributed rate prior (9-10). Posterior estimates of molecular substitution rates and divergence times were estimated during 200,000 generations of MCMC, and results inspected for stationarity in the program Tracer (11). To overcome computational limitations with genome scale data we subset 15 loci for use in our MCMCtree analysis using principal component axis 2 in the gensortR (see ref. 12 for details).

**Evolution of life history and edaphic endemism**

All analyses were implemented in RevBayes (12) using the post burn-in MCC chronogram output of our MCMCtree analysis with outgroup taxa removed. Maximum a posteriori (MAP) estimates of ancestral states for both life history and edaphic endemism were based on a post burn-in posterior of 13,500 trees. We used reversible jump MCMC to test for reversibility by alternatively setting each transition rate to zero or inferring non-zero transition rates under an exponentially distributed prior. Based on past sensitivity analyses (2), non-zero transition rates were drawn from an exponential distribution with a mean of 1 transition over the tree. Reversible or irreversible models were inferred with equal prior probability and posterior model estimates recorded using a custom binary logical monitor. We used Bayes Factors to assess support for irreversibility by calculating the natural log of the number of iterations in which an irreversible model was inferred (the posterior odds of model 0) divided by the number of iterations where a reversible model was inferred (the posterior odds of model 1), while disregarding the prior odds since we used a uniform prior. Given proven sensitivity of tests for reversibility to prior information from root states (14), we based root state frequencies on ancestral state reconstructions of habitat and life-history derived from a previous study of helenioid Heliantheae (15), which showed strong prior support for tropical and perennial states at the root of Perityleae. Transition rates under all combinations of irreversible and reversible models were inferred during 15,000 iterations of an MCMC and inspected for convergence. To improve mixing, in the absence of support for irreversibility, the analysis was rerun under the non-zero transition rate prior and (MAP) ancestral states compiled from a post burn-in posterior of 13,500 trees.

To test for associations between life history and endemism on rocky outcrops we used BayesTraits (16) to compare the fit (log-likelihood) of two continuous time Markov models of life history and edaphic evolution, a dependent and independent model (17). Both models aimed to infer transitions to or from the suffrutescent perennial life-history and bare rock edaphic affinity at any given moment over infinitesimally small units of time, but the dependent model included an additional four transition-rate parameters (eight total) because it assumes that the rate of change in one trait is dependent on the state of the other. Double transitions in both traits at the same time were disallowed in both models. Given the size of the data and complexity of the models, we used reversible jump MCMC (rjMCMC) to integrate parameter estimates over the model space, weighted by their probability. Models were compared using log Bayes Factors with the equation: Log BF = 2(log likelihood dependent model – log likelihood independent model) and a log BF greater than 10 was interpreted as strong support for a correlation. For each model rjMCMC was conditioned on an exponential distribution with a mean of 100 and a hyperprior of zero. Marginal log-likelihoods were calculated using a steppingstone sampler (18) with 100 stones successively heated for 10,000 iterations.

**Historical biogeography with paleo-biome informed models**

Discrete biogeographical ranges for each taxon were obtained from the Southwestern Environmental Information Network (SEInet) and from an extensive review of herbarium specimens as part of the first author's systematic research on Perityleae. We scored each tip for presence in four geographical regions: South America, Baja California, the Trans-Mexican Volcanic Belt, and the Basin and Range Province (Fig. 2, S4). In South America, Perityleae has a disjunct distribution, occurring in the Atacama Desert in southern Peru and northern Chile as well as the Desventuradas Islands, ~200 km off the coast of central Chile. Baja California was defined as including the entire peninsula and nearby islands, including Guadalupe Island, the Channel Islands of southwest California, and the Revillagigedo Archipelago, located 100 km south of San Jose del Cabo. The Trans-Mexican Volcanic Belt (19) described the area covered by the predominantly volcanic mountain ranges and plateaus of northwest and central Mexico, including the Sierra Madre Occidental. The Basin and Range Province covers a wide geographic area of western North America that is characterized by its distinctive topography, with thousands of discontinuous mountain ranges punctuated by low, arid valleys (19). The Basin and Range Province here included the Madrean sky island archipelago and surrounding Sonoran Desert, the eastern continental Chihuahuan Desert and sky islands of Texas and New Mexico in the United States, and Chihuahua, Coahuila, and Tamaulipas in Mexico, as well as the numerous north-south oriented mountain ranges of the Great Basin region in Nevada and Utah, which have been called "caterpillars marching toward Mexico." (20)

Our characterization of biomes was based on the level I eco-regions of North America published by the Intergovernmental Commission for Environmental Cooperation. Desert includes the Great Basin, Sonoran, and Chihuahuan subregions, which cover broad swaths of the arid southwest United States, Baja California, and northern mainland Mexico. This biome includes both hot and cold deserts with unpredictable precipitation averaging ~250 mm or less per year and arid plant communities dominated by shrubs, stem succulents, and seasonal annuals scattered in arid, mostly open landscapes without a dense canopy cover (19). The temperate biome included high elevations of the Sierra Madre Occidental in Mexico and the Mogollon Rim in Arizona, the only areas in our study region where precipitation is predictably high, yet temperatures fall seasonally below freezing, containing plant communities dominated by coniferous or deciduous trees (19). The subtropical biome includes mid-elevation parts of the arid southwest U.S. and several states in northern and central Mexico where periodic but not extreme drought combined with infrequent freezing temperatures produces a vegetation mosaic of grassland and shrubland with patches of forest (19). The tropical biome mostly refers to tropical deciduous forests (selva tropical subcaudicifolio sensu 19), habitats where freezing temperatures do not regularly occur and seasonality in rainfall is intense, and which support a dense canopy of mostly drought deciduous trees, robust multi-stemmed shrubs, and stem succulents. In our study area, tropical deciduous forests cover the western Pacific plain of the Sierra Madre Occidental as well as the southern Cape Region and Sierra la Giganta of Baja California (19, 21-22). For a majority of Perityleae, we were able to assign each taxon to only one biome. The exceptions were wide ranging taxa, which were coded as polymorphic for multiple biomes with equal probability. One species, *Perityle emoryi*, has a distribution that overlaps with the California Floristic Province, but is mostly found in desert habitats, so we assigned it to North American desert (23). *Nesothamnus incanus,* which is endemic to the environmentally heterogeneous Guadalupe Island off the coast of Baja California, Mexico is found in low, arid habitats within the island and was therefore assigned to desert (21).

Prior to running a fully parameterized biome shift model, we first estimated rates of dispersal and extirpation among geographic areas, as well as ancestral biogeographic ranges separately from biomes using a Bayesian implementation of the Dispersal Extinction Cladogenesis (DEC) model (24) in RevBayes (13). We based our analyses on the MAP chronogram output of our MCMCtree analysis. Given evidence that transitions into the null range can result in abnormal extirpation rates and range sizes, we conditioned our analysis on survival by setting the probability of entering the null range to zero (25). Our implementation of the DEC modeled anagenetic dispersal and local extinction (extirpation) rates based a log-uniform prior and considered only allopatry and subset sympatry as potential cladogenetic events (see 26 for rationale). Dispersal and extirpation rates, cladogenetic events, and ancestral biogeographic ranges were inferred during 30,000 iterations of an MCMC. MAP estimates of ancestral ranges at nodes and shoulders were compiled from 28,000 samples of post burn-in posterior trees and the results plotted using the R package RevGadgets (27).

To explicitly test the hypothesis that shifts into the desert biome tracked the availability of desert habitats through time, we modeled two different scenarios: a) total dependence of biome-shifts on paleo-biome structure through time and b) total independence of shifts on paleo-biome structure through time. To model independence we used the set.value() function in RevBayes to assign zero as a prior on factor w_b and 0.5 to factors w_u and w_g, and to model dependence w_b was assigned a value of 1 and w_u and w_g a value of 0. We then estimated transition rates among biomes and generated model averaged ancestral states across the four epochs during two separate iterations of 25,000 generations of MCMC. Each run was checked for convergence in the program Tracer and marginal likelihoods calculated using thermodynamic integration with stepping stone sampling at a tuning interval of 50 for 1000 generations. Log bayes factors were calculated from the marginal likelihoods as Log BF = 2(log likelihood dependent model – log likelihood independent model) and a log BF of greater than 10 interpreted as strong support for a dependent model.

**Supplemental results and discussion**

**Taxonomic implications of a more densely sampled phylogeny of Perityleae**

Unambiguous resolution of a diverse early diverging lineage that tracked the separation of the Baja California Peninsula reinforces the recently improved understanding of Perityleae that required revision of taxonomic concepts (28). Until recently, most of the species diversity in Perityleae was recognized within the "core" genus *Perityle,* but non-monophyly of *Perityle* and support for the type species, *Perityle californica*, as nested within an early diverging Baja California clade was the basis for several changes, including: a narrowed circumscription of *Perityle,* recognition of the long-branch *Nesothamnus,* and reinstatement of *Galinsogeopsis* and *Laphamia* with new, broader delimitations. Expanded sampling of the Baja California clade using target capture in the current study also supports the inclusion within *Perityle* of *P. socorrensis* Rose*, P. emoryi* Torr.*, Perityle tenuifolius* (Phil.) Lichter-Marck, and the three taxa of genus *Amauria* Benth. (22, 28). *Perityle* remains non-monophyletic as currently circumscribed, with

*Perityle rosei* Greenm. and *P. trichodonta* S.F. Blake resolved as more closely related to *Pericome* and *Eutetras* than to other members of *Perityle.* The monotypic, Guadalupe Island endemic *Nesothamnus incanus* (A. Gray) Rydb. occupies a long branch that is the sister lineage to *Galinsogeopsis*.

**Non-monophyly of densely sampled taxa**

Inclusion of multiple conspecific samples in the present study showed support for monophyly of recognized species in most cases, but there were several species for which samples did not group together. Taxa found to be non-monophyletic include *Galinsogeopsis feddemae* (McVaugh) I.H. Lichter-Marck, *Laphamia cordifolia* (S.F. Blake) I.H. Lichter-Marck, *L. gracilis* M.E. Jones, *L. lemmonii* A. Gray, *L. megalocephala* S. Watson, *L. rupestris* A. Gray, *Perityle cuneata*, *P. emoryi, and P. rosei*. In part, these results reflect our choice to sequence samples representative of morphologically unique and geographically disjunct populations discovered during the first author's field work and examination of herbarium collections. Perityleae tend to grow in unreachable parts of the most inaccessible cliffs and canyons in the North American deserts, where their isolated populations are arrayed across 2000 miles of dry and mostly roadless terrain punctuated by imposing rocky mountains (28). Comprehensive floristic studies of sky islands and parts of the Sierra Madre Occidental have led to the description of many new-to-science species of Perityleae since the tribe was last treated by Powell (e.g., 29), and our results suggest that the currently recognized diversity of Perityleae continues to be underestimated. Our fragmentary knowledge of the diversity in this group is a conservation concern because Perityleae habitats are imminently threatened by energy development, climate change, and mining. With >30% of the species in this group listed as vulnerable, imperiled, or critically endangered on NatureServe, Perityleae stands out as having one of the highest proportions of at-risk taxa of any taxon in the arid western North American flora.

**Widespread occurrence of edaphic endemism to bare habitats among close relatives to predominantly desert plant and animal lineages**

Familiar desert plant lineages are dominant elements in the flora of rocky outcrops in the tropical deciduous forests that are adjacent to, and considered a major source area for expansions into, the North American deserts (31), including cacti (Cactaceae; e.g. *Echinocereus bacanorensis* (W. Rischer & Trocha) W. Blum and *Mammillaria grahamii* Engelm.), agaves (Agavaceae; e.g. *Agave aurea* Brandegee), euphorbias (Euphorbiaceae; e.g. *Euphorbia chaetocalyx* var. *triligulata* (L.C. Wheeler). M.C. Johnst.), legumes (Leguminosae; e.g. *Erythrina flabelliformis* Kearney), cheilanthoid ferns (Pteridaceae; e.g. *Astrolepis sinuata* Sw., *Notholaena aurantiaca* D.C. Eaton, *Myriopteris myriophylla* (Desv.) J.Sm., and *Argyrochosma peninsularis* (Maxon & Weath.) Windham), composites (Compositae; e.g. *Brickellia brandegeei* B.L. Rob., *Coreocarpus dissectus* S.F. Blake, and *Xanthisma incisifolium* (I.M. Johnst.) G. Nesom), evening primroses (Onagraceae; e.g. *Oenothera toumeyi* Tidestr. and *Gongylocarpus fruticulosus* Brandegee), mints (Lamiaceae; *Hedeoma drummundii* Benth.), stonecrops (Crassulaceae; e.g. *Graptopetalum rusbyi* Rose, *Echeverria colorata* E. Walther, and *Dudleya nubigena* Britton & Rose), chuparosas (Acanthaceae; e.g. *Justicia sonorae* Wassh.), turpentine-brooms (Rutaceae; e.g. *Thamnosma ciliata* I.M. Johnst.), and datillos (Bromeliaceae; e.g., *Hechtia* jaliscana L. B. Sm.). Decomposed, but typically open edaphic substrates such as gypsum, greenstone, and limestone outcrops, alkaline sinks, and volcanic talus also harbor members or close relatives of predominantly desert plant lineages in adjacent more densely vegetated biomes, including *Tiquilia* (Ehretiaceae; e.g. *T. gossypina* clade), *Encelia* (Compositae; e.g. *Enceliopsis* spp.), ivesioids (Rosaceae; *Potentilla* spp.), jewelflowers (Brassicaceae; Thelypodieae), *Nama* (Namaceae; e.g. *Nama rupicola* Bonpl. ex Choisy), and *Zeltnera* (Gentianaceae; e.g. *Zeltnera calycosa* (Buckley) G. Mans.). Similar patterns can be found in groups of predominantly desert animals, such as rattlesnakes, lizards, geckos, and tortoises, which are often found closely associated with rocky outcrops in a tropical context, though arboreal lifestyles may have also played a role in pre-adapting these lineages for success in the desert ecosystem (30).

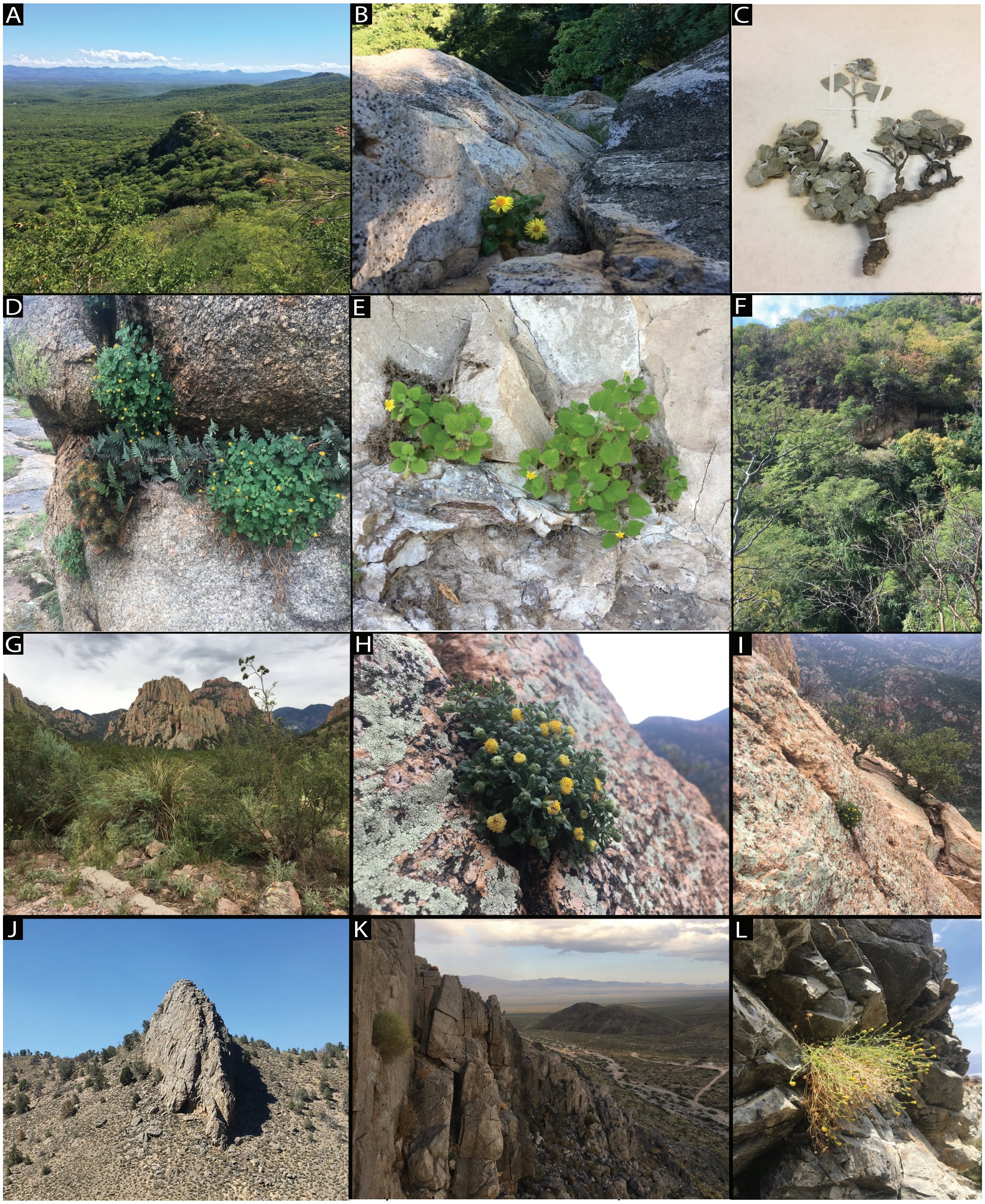

**Fig. S1.** Additional photographs illustrating habit and habitat diversity within the predominantly desert and tropical deciduous forest dwelling rock daisy genus *Laphamia*. (A) A scattered rocky outcrop within otherwise densely vegetated tropical deciduous forest in the Sierra de Alamos, Sonora, Mexico. (B) Growth out of the exposed cliffs on the eastern side of the outcrop shown in (A) of *Laphamia cordifolia*. (C) An herbarium specimen showing the perennating woody caudex possessed by suffrutescent perennial Perityleae that is associated with growth on bare, rocky outcrops. (D) A rock outcrop plant community in a tropical deciduous forest in Sonora, Mexico including members of the following otherwise predominantly desert plant groups: cacti

(Cactaceae, *Echinocereus bacanorensis*), cheilanthoid ferns (Cheilanthoideae, *Myriopteris lindheimeri),* and rock daisies (Perityleae, *Laphamia reinana*). ( E) *Laphamia gentryi* growing on a rocky outcrop surrounded by (F) tropical deciduous forest in the western Sierra Madre Occidental in Sonora, Mexico. (G) Cliffs and cathedrals of rhyolite in the Chiricahua Mountains in Arizona, U.S.A. (H) *Laphamia cochisensis,* a narrow endemic to the Chiricahua Mountains growing out of bare, rhyolitic cliffs. (I) View of surrounding subtropical forest from the vantage point of *Laphamia cochisensis.* (J) Limestone outcrops of the Great Basin Desert in the Inyo Mountains, California, U.S.A. (K) *Laphamia intricata* (left foreground) growing on calcareous outcrops in the Ivanpah Mountains, California, U.S.A. (L) *Laphamia villosa*, a narrow endemic found only on limestone outcrops in the Panamint Range, California, U.S.A.

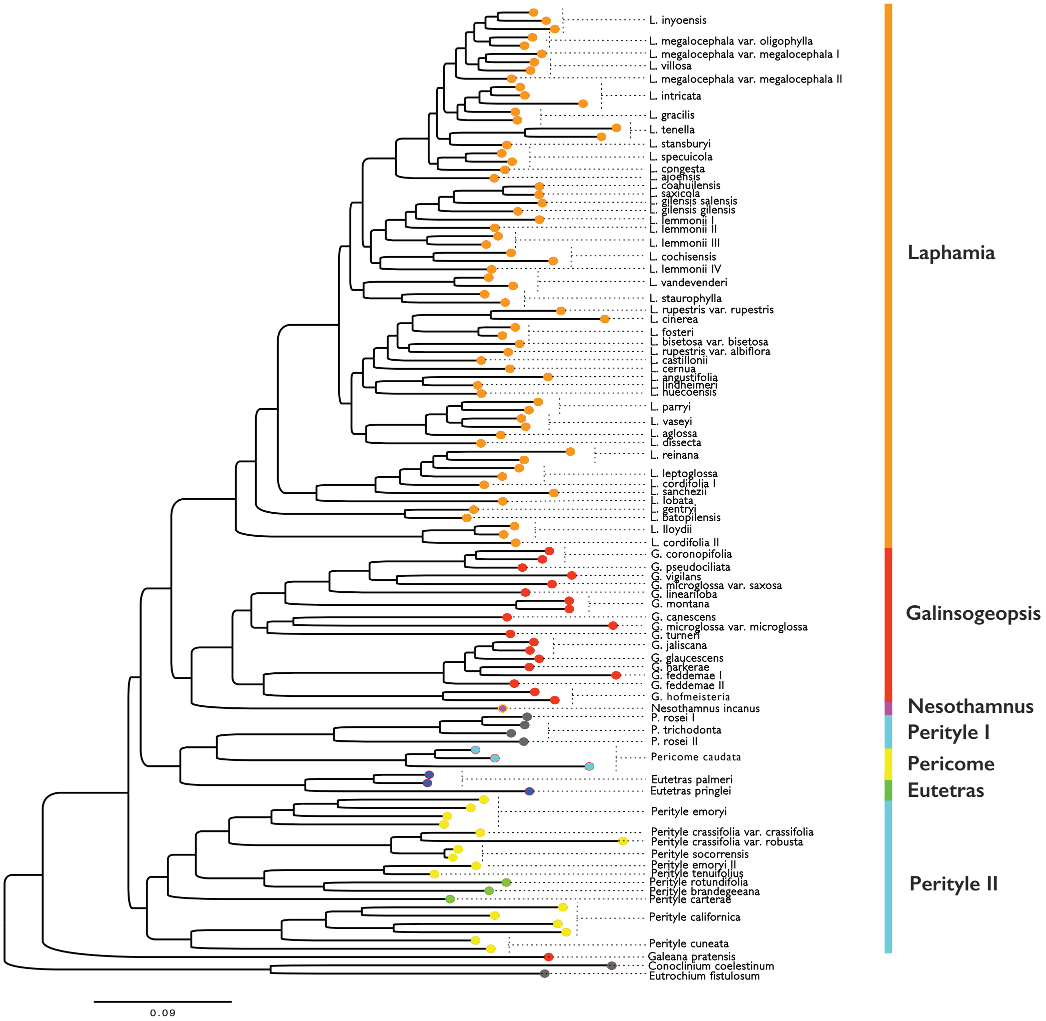

**Fig. S2.** Phylogeny of Perityleae based on concatenated analysis of 229 low-copy nuclear supercontigs combining both exons and introns using Bayesian statistical inference in the program ExaBayes (7). The data matrix on which this analysis was based was ~1,400,000 bp. Posterior probabilities were estimated during 1,000,000 iterations of MCMC and checked for stationarity in the program Tracer. Posterior probabilities of 1 were recovered at all nodes. Color coded tips denote genera in the recent classification by Lichter-Marck (28). The outgroup *Helianthus annuus* not shown.

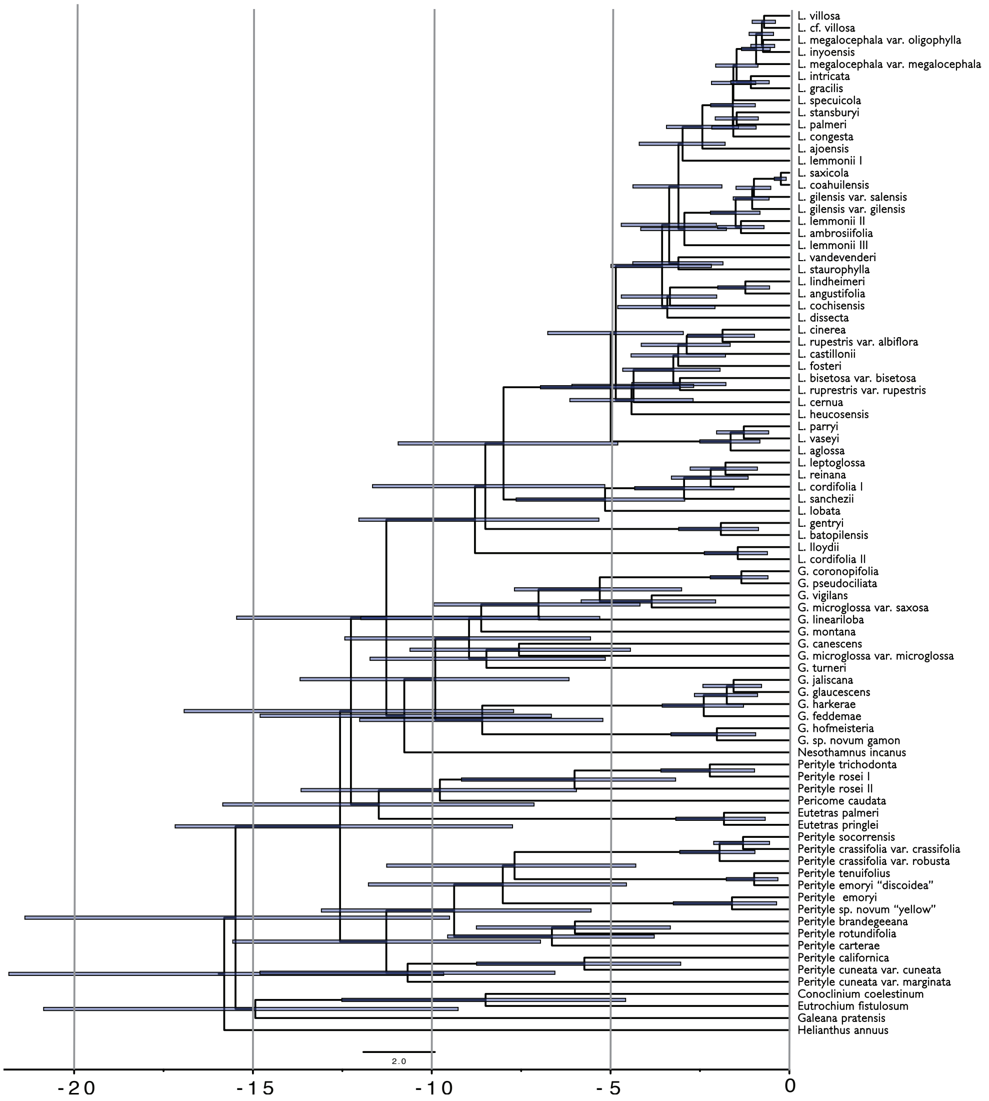

**Fig. S3.** Time calibrated phylogeny of Perityleae based on fixed topology derived from a pseudocoalescent analysis of 229 low-copy nuclear genes, with branch lengths inferred using the approximation method in MCMCtree using a 15 loci obtained using phylogenetic subsampling (9-10). Our analysis was conditioned on an uncorrelated log normal relaxed molecular clock and an HKY molecular substitution model with four gamma distributed rate categories. Posterior estimates of divergence times were obtained during 50,000 iterations of MCMC and the analysis repeated to check for comparable results. Scale bars: 2 my.

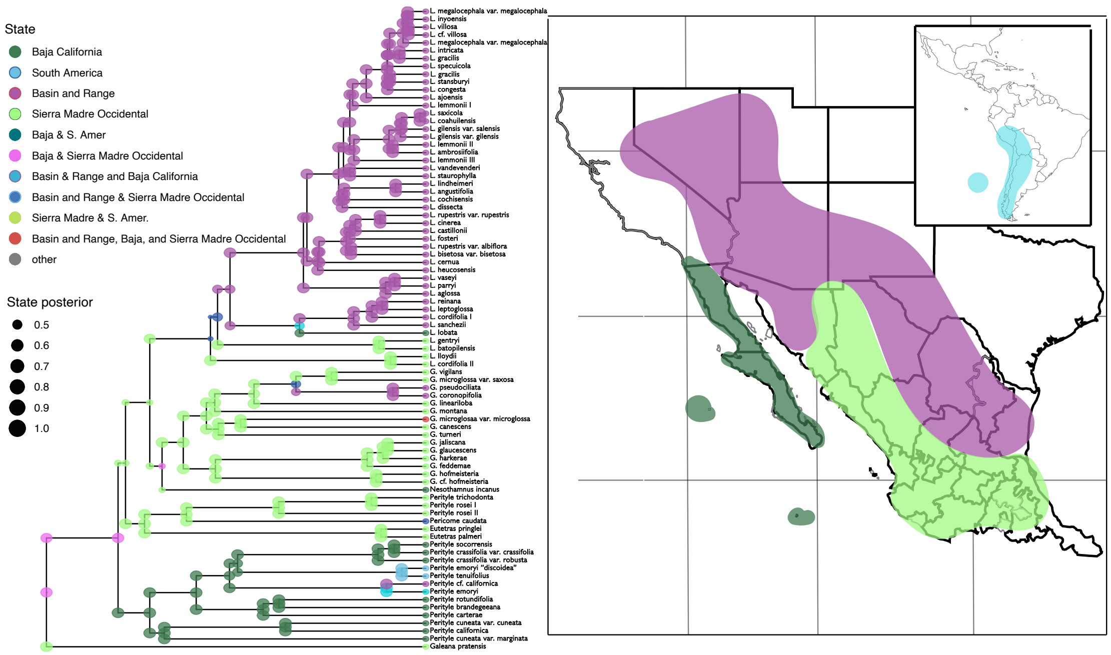

**Fig. S4.** Maximum a posteriori ancestral state reconstruction of biogeographic ranges of Perityleae at nodes and shoulders based on a dispersal-extinction-cladogenesis model implemented in RevBayes. Colors correspond to the geographic regions illustrated on the right and colored node-dot size is scaled to posterior probability. Cladogenetic events are indicated as instances where the daughter ranges differ from the common ancestor's range at their intervening node.

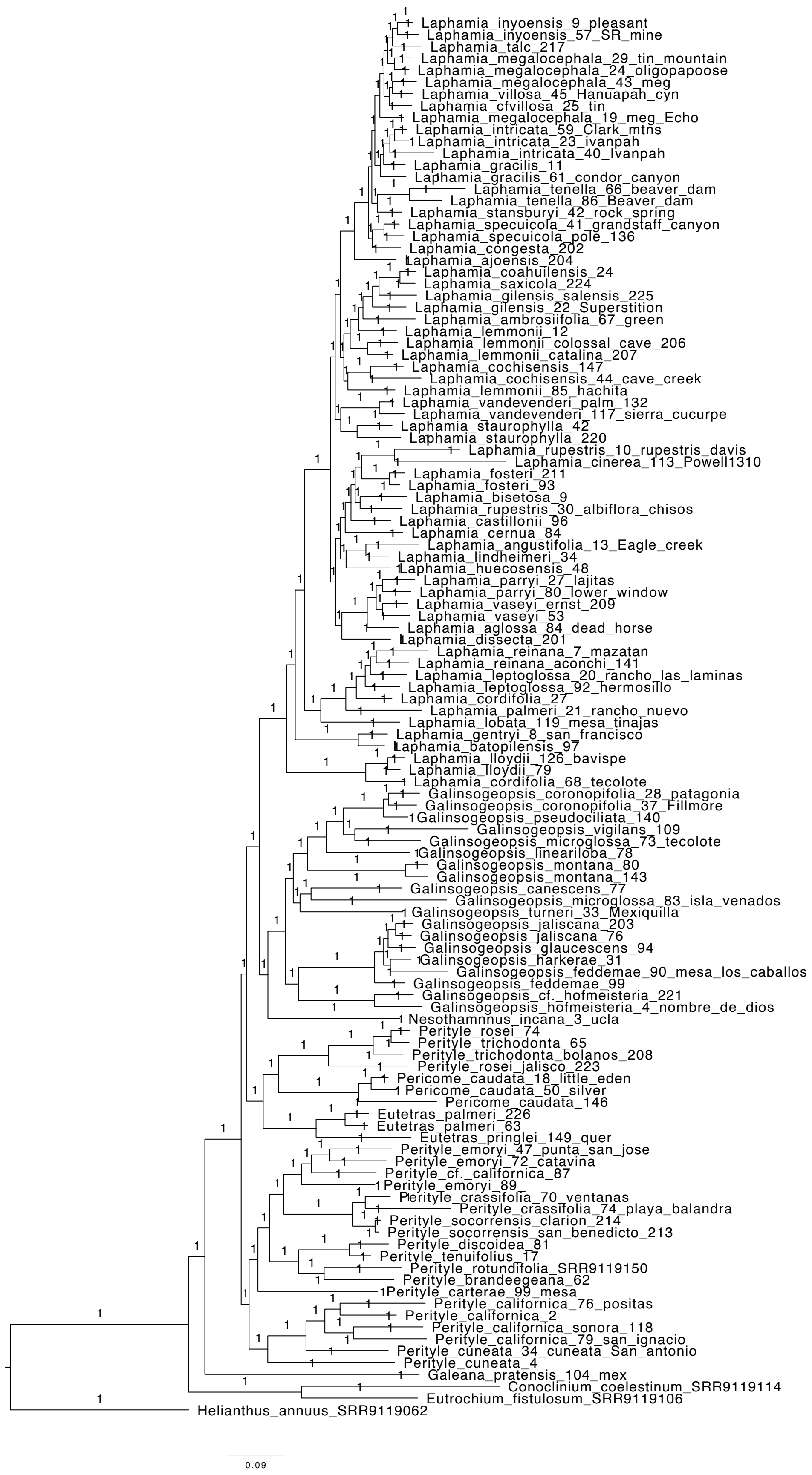

Laphamia_inyoensis_9_pleasant
Laphamia_inyoensis_57_SR_mine
Laphamia_talc_217
Laphamia_megalocephala_29_tin_mountain
Laphamia_megalocephala_24_oligopapoose
Laphamia_megalocephala_43_meg
Laphamia_villosa_45_Hanuapah_cyn
Laphamia_cfvillosa_25_tin
Laphamia_megalocephala_19_meg_Echo
Laphamia_intricata_59_Clark_mtns
Laphamia_intricata_23_ivanpah
Laphamia_intricata_40_Ivanpah
Laphamia_gracilis_11
Laphamia_gracilis_61_condor_canyon
Laphamia_tenella_66_beaver_dam
Laphamia_tenella_86_Beaver_dam
Laphamia_stansburyi_42_rock_spring
Laphamia_specuicola_41_grandstaff_canyon
Laphamia_specuicola_pole_136
Laphamia_congesta_202
Laphamia_ajoensis_204
Laphamia_coahuilensis_24
Laphamia_saxicola_224
Laphamia_gilensis_salensis_225
Laphamia_gilensis_22_Superstition
Laphamia_ambrosiifolia_67_green
Laphamia_lemmonii_12
Laphamia_lemmonii_colossal_cave_206
Laphamia_lemmonii_catalina_207
Laphamia_cochisensis_147
Laphamia_cochisensis_44_cave_creek
Laphamia_lemmonii_85_hachita
Laphamia_vandevenderi_palm_132
Laphamia_vandevenderi_117_sierra_cucurpe
Laphamia_staurophylla_42
Laphamia_staurophylla_220
Laphamia_rupestris_10_rupestris_davis
Laphamia_cinerea_113_Powell1310
Laphamia_fosteri_211
Laphamia_fosteri_93
Laphamia_bisetosa_9
Laphamia_rupestris_30_albiflora_chisos
Laphamia_castillonii_96
Laphamia_cernua_84
Laphamia_angustifolia_13_Eagle_creek
Laphamia_lindheimeri_34
Laphamia_huecosensis_48
Laphamia_parryi_27_lajitas
Laphamia_parryi_80_lower_window
Laphamia_vaseyi_ernst_209
Laphamia_vaseyi_53
Laphamia_aglossa_84_dead_horse
Laphamia_dissecta_201
Laphamia_reinana_7_mazatan
Laphamia_reinana_aconchi_141
Laphamia_leptoglossa_20_rancho_las_laminas
Laphamia_leptoglossa_92_hermosillo
Laphamia_cordifolia_27
Laphamia_palmeri_21_rancho_nuevo
Laphamia_lobata_119_mesa_tinajas
Laphamia_gentryi_8_san_francisco
Laphamia_batopilensis_97
Laphamia_lloydii_126_bavispe
Laphamia_lloydii_79
Laphamia_cordifolia_68_tecolote
Galinsogeopsis_coronopifolia_28_patagonia
Galinsogeopsis_coronopifolia_37_Fillmore
Galinsogeopsis_pseudociliata_140
Galinsogeopsis_vigilans_109
Galinsogeopsis_microglossa_73_tecolote
Galinsogeopsis_lineariloba_78
Galinsogeopsis_montana_80
Galinsogeopsis_montana_143
Galinsogeopsis_canescens_77
Galinsogeopsis_microglossa_83_isla_venados
Galinsogeopsis_turneri_33_Mexiquilla
Galinsogeopsis_jaliscana_203
Galinsogeopsis_jaliscana_76
Galinsogeopsis_glaucescens_94
Galinsogeopsis_harkerae_31
Galinsogeopsis_feddemae_90_mesa_los_caballos
Galinsogeopsis_feddemae_99
Galinsogeopsis_cf._hofmeisteria_221
Galinsogeopsis_hofmeisteria_4_nombre_de_dios
Nesothamnnus_incana_3_ucla
Perityle_rosei_74
Perityle_trichodonta_65
Perityle_trichodonta_bolanos_208
Perityle_rosei_jalisco_223
Pericome_caudata_18_little_eden
Pericome_caudata_50_silver
Pericome_caudata_146
Eutetras_palmeri_226
Eutetras_palmeri_63
Eutetras_pringlei_149_quer
Perityle_emoryi_47_punta_san_jose
Perityle_emoryi_72_catavina
Perityle_cf._californica_87
Perityle_emoryi_89_
Perityle_crassifolia_70_ventanas
Perityle_crassifolia_74_playa_balandra
Perityle_socorrensis_clarion_214
Perityle_socorrensis_san_benedicto_213
Perityle_discoidea_81
Perityle_tenuifolius_17
Perityle_rotundifolia_SRR9119150
Perityle_brandeegeana_62
Perityle_carterae_99_mesa
Perityle_californica_76_positas
Perityle_californica_2
Perityle_californica_sonora_118
Perityle_californica_79_san_ignacio
Perityle_cuneata_34_cuneata_San_antonio
Perityle_cuneata_4
Galeana_pratensis_104_mex
Conoclinium_coelestinum_SRR9119114
Eutrochium_fistulosum_SRR9119106
Helianthus_annuus_SRR9062
0.09

**Fig. S5.** Maximum clade credibility phylogeny of Perityleae based on a partitioned data matrix of 229 low-copy nuclear loci inferred using Bayesian statistical inference in ExaBayes (7). Posterior probabilities were estimated during 1,000,000 generations of MCMC.

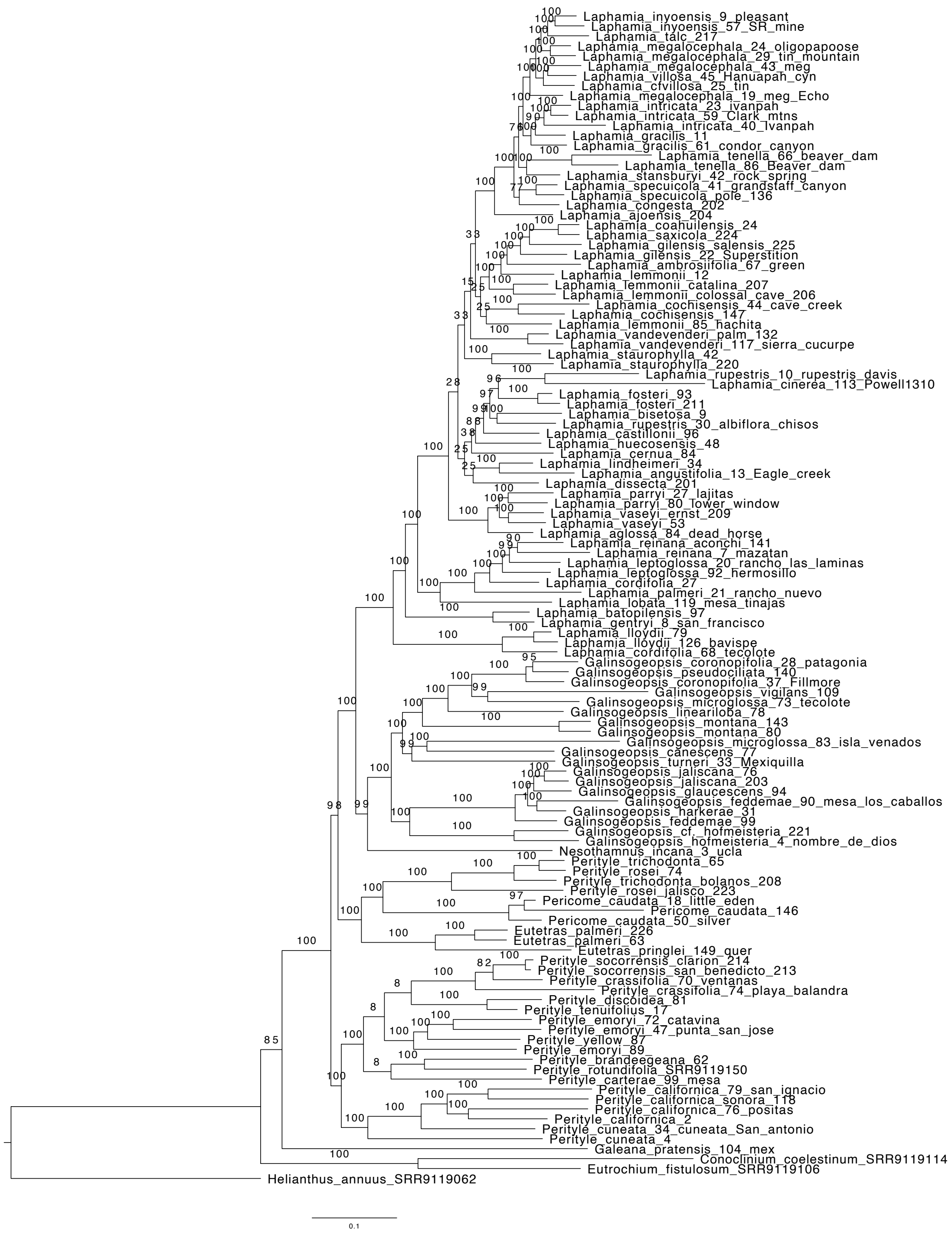

**Fig. S6.** Maximum likelihood phylogeny of Perityleae based on a partitioned data matrix of 229 low-copy nuclear genes inferred using a GTR+gamma+I molecular substitution model in RAxML-NG.

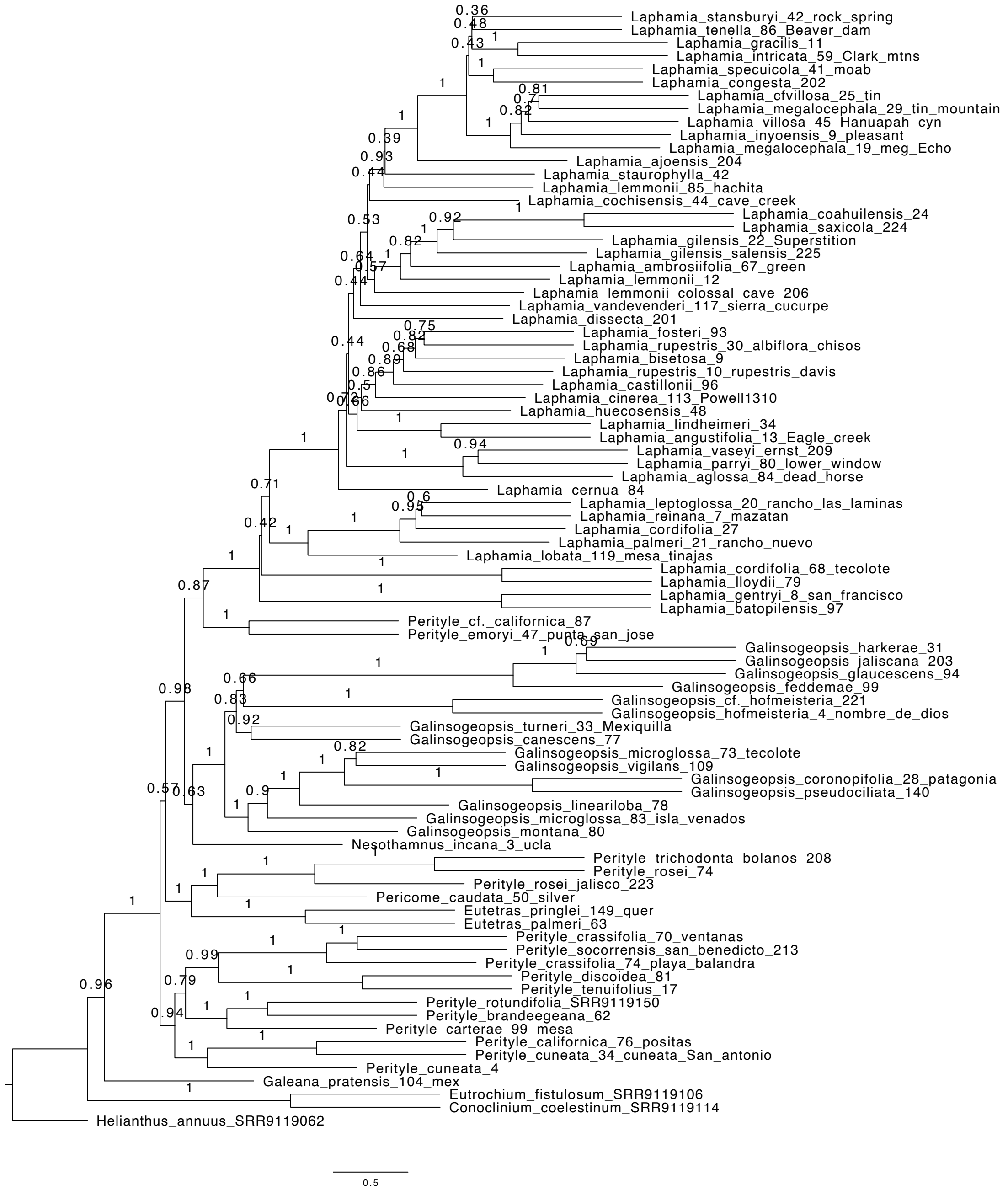

**Fig. S7.** Pseudo-coalescent phylogeny of Perityleae based on ASTRAL-III analysis of 229 gene trees inferred separately in RAxML under a GTRCAT substitution model.

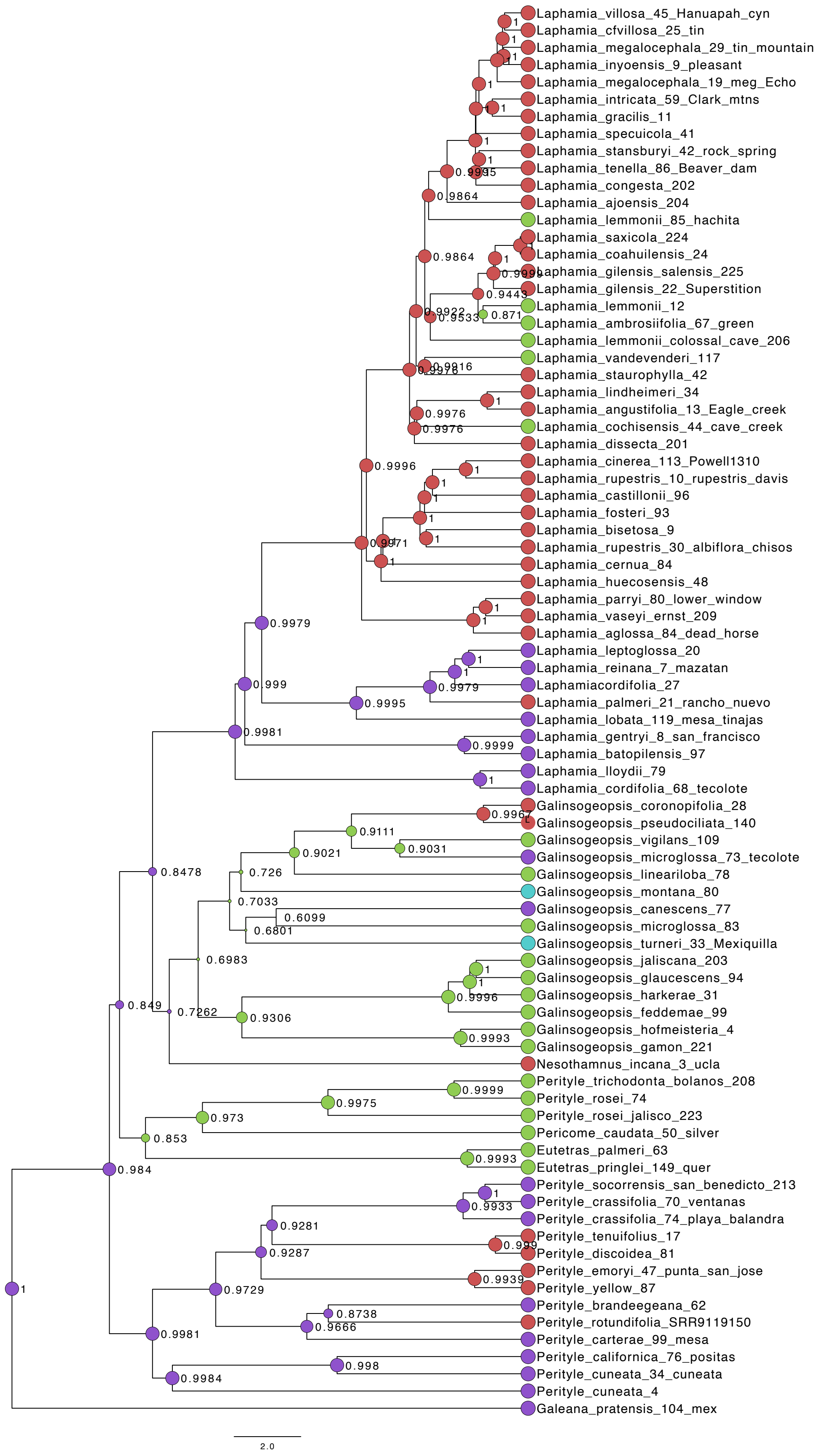
Laphamia_villosa_45_Hanuapah_cyn
Laphamia_cfvillosa_25_tin
Laphamia_megalocephala_29_tin_mountain
Laphamia_inyoensis_9_pleasant
Laphamia_megalocephala_19_meg_Echo
Laphamia_intricata_59_Clark_mtns
Laphamia_gracilis_11
Laphamia_specuicola_41
Laphamia_stansburyi_42_rock_spring
Laphamia_tenella_86_Beaver_dam
Laphamia_congesta_202
Laphamia_ajoensis_204
Laphamia_lemmonii_85_hachita
Laphamia_saxicola_224
Laphamia_coahuilensis_24
Laphamia_gilensis_salensis_225
Laphamia_gilensis_22_Superstition
Laphamia_lemmonii_12
Laphamia_ambrosiifolia_67_green
Laphamia_lemmonii_colossal_cave_206
Laphamia_vandevenderi_117
Laphamia_staurophylla_42
Laphamia_lindheimeri_34
Laphamia_angustifolia_13_Eagle_creek
Laphamia_cochisensis_44_cave_creek
Laphamia_dissecta_201
Laphamia_cinerea_113_Powell1310
Laphamia_rupestris_10_rupestris_davis
Laphamia_castillonii_96
Laphamia_fosteri_93
Laphamia_bisetosa_9
Laphamia_rupestris_30_albiflora_chisos
Laphamia_cernua_84
Laphamia_huecosensis_48
Laphamia_parryi_80_lower_window
Laphamia_vaseyi_ernst_209
Laphamia_aglossa_84_dead_horse
Laphamia_leptoglossa_20
Laphamia_reinana_7_mazatan
Laphamiacordifolia_27
Laphamia_palmeri_21_rancho_nuevo
Laphamia_lobata_119_mesa_tinajas
Laphamia_gentryi_8_san_francisco
Laphamia_batopilensis_97
Laphamia_lloydii_79
Laphamia_cordifolia_68_tecolote
Galinsogeopsis_coronopifolia_28
Galinsogeopsis_pseudociliata_140
Galinsogeopsis_vigilans_109
Galinsogeopsis_microglossa_73_tecolote
Galinsogeopsis_leariloba_78
Galinsogeopsis_montana_80
Galinsogeopsis_canescens_77
Galinsogeopsis_microglossa_83
Galinsogeopsis_turneri_33_Mexiquilla
Galinsogeopsis_jaliscana_203
Galinsogeopsis_glaucescens_94
Galinsogeopsis_harkerae_31
Galinsogeopsis_feddemae_99
Galinsogeopsis_hofmeisteria_4
Galinsogeopsis_gamon_221
Nesothamnus_incana_3_ucla
Perityle_trichodonta_bolanos_208
Perityle_rosei_74
Perityle_rosei_jalisco_223
Pericome_caudata_50_silver
Eutetras_palmeri_63
Eutetras_pringlei_149_quer
Perityle_socorrensis_san_benedicto_213
Perityle_crassifolia_70_ventanas
Perityle_crassifolia_74_playa_balandra
Perityle_tenuifolius_17
Perityle_discoidea_81
Perityle_emoryi_47_punta_san_jose
Perityle_yellow_87
Perityle_brandeegeana_62
Perityle_rotundifolia_SRR9119150
Perityle_carterae_99_mesa
Perityle_californica_76_positas
Perityle_cuneata_34_cuneata
Perityle_cuneata_4
Galeana_pratensis_104_mex
2.0

**Fig. S8.** Maximum a posteriori reconstruction of ancestral habitat in Perityleae estimated using MCMC in RevBayes. Purple = tropical, green = subtropical, blue = temperate, and red = desert. Values at nodes correspond to posterior probabilities.

**Table S1.** Voucher information, and GenBank SRA accession numbers for sampled individuals of Perityleae sequenced for this study. See ref. 2 for information on samples included in this previous study. Herbarium acronyms follow Thiers, B. [continuously updated]. Index Herbariorum: A global directory of public herbaria and associated staff. New York Botanical Garden's Virtual Herbarium. http://sweetgum.nybg.org/science/ih/

| Taxon | Accession or collection # | Location | Herbarium | Collector(s) | Collection date | GenBank SRA # |
|---|---|---|---|---|---|---|
| *Laphamia inyoensis* Ferris | ILM 48 | Pleasant Mountain, Inyo Co. CA, USA | UC | I.H. Lichter-Marck & K. Allerton | July 2018 | |
| *Laphamia* cf. *inyoensis* Ferris | ILM 105 | Talc City, Inyo Co., CA, USA | UC | I.H. Lichter-Marck & K. Allerton | July 2018 | |
| *Laphamia megalocephala* var. *oligophylla* (A.M. Powell) I.H. Lichter-Marck | ILM 204 | Tin Mountain, Inyo Co., CA, USA | UC | I.H. Lichter-Marck & K. Allerton | July 2018 | |
| *Laphamia megalocephala* var. *oligophylla* (A.M. Powell) I.H. Lichter-Marck | ILM 166 | Papoose Flat, Inyo Co., CA, USA | UC | I.H. Lichter-Marck & K. Allerton | July 2018 | |
| *Laphamia megalocephala* var. *megalocephala* (S. Watson) | ILM 334 | Echo Canyon Reservoir, Pioche, NV, USA | UC | I.H. Lichter-Marck | September 2018 | |
| *Laphamia intricata* Brandegee | ILM 229 | Ivanpah Mountains, San Bernardino Co., CA, USA | UC | I.H. Lichter-Marck | August 2018 | |
| *Laphamia intricata* Brandegee | ILM 230 | Ivanpah Mountains, San Bernardino Co., CA, USA | UC | I.H. Lichter-Marck | August 2018 | |
| *Laphamia gracilis* M.E. Jones | UC 1582885 | Sheep Range, Clark Co., NV, USA | UC | T.L. Ackerman | 2 July 1979 | |
| *Laphamia palmeri* A. Gray | ILM 339 | Beaver Dam Mountains, Washington Co., UT, USA | UC | I.H. Lichter-Marck | September 2018 | |
| *Laphamia specuicola* (S.L. Welsh | ILM 322 | Moab, Grand Co., UT, USA | UC | I.H. Lichter-Marck | September 2018 | |

| | | | | | |
|---|---|---|---|---|---|
| & Neese) I.H. Lichter-Marck | | | | | |
| *Laphamia congesta* M.E. Jones | ILM 874 | South Rim, Grand Canyon, Coconino Co., Arizona, USA | UC | I.H Lichter-Marck & S. Winitsky | Septemb er 2019 |
| *Laphamia ajoensis* (Todsen) I.H Lichter-Marck | ILM | Ajo Mountains, Pima Co., AZ, USA | UC | I.H Lichter-Marck & S. Winitsky | Decembe r 2020 |
| *Laphamia coahuilensus* (A.M. Powell) I.H. Lichter-Marck | AZ 337203 | Sierra de San Marcos, Coahuila, MX | AZ | M.C. Johnston, T. Wendt, & F. Chiang. | 12 June 1972 |
| *Laphamia saxicola* Eastw. | AZ 372727 | Superstition Mountains, Maricopa Co., AZ, USA | AZ | M. Quinn | 17 April 2003 |
| *Laphamia gilensis* var. *salensis* (A.M. Powell) I.H. Lichter-Marck | ILM 314 | Salt River, Apache Co., AZ, USA | UC | I.H. Lichter-Marck | August 2018 |
| *Laphamia gilensis* var. *gilensis* (M.E. Jones) | UCR 138793 | Tortilla Flat, Superstition Mountains, AZ, USA | UC | W. Hodgson | 15 April 1993 |
| *Laphamia lemmonii* A. Gray | UC 709590 | Graham Mountains, Graham Co., AZ, USA | UC | R.M. McGuire. | 2 June 1935 |
| *Laphamia lemmonii* A. Gray | ILM 310 | Colossal Cave, Pima Co., AZ, USA | UC | I.H. Lichter-Marck | August 2018 |
| *Laphamia lemmonii* A. Gray | ILM 274 | Catalina Mountains, Pima Co., AZ, USA | UC | I.H. Lichter-Marck & S. Winitsky | Decembe r 2020 |
| *Laphamia cochisensis* W.E. Niles | ILM 789 | Chiricahua Mountains, Cochise Co., AZ, USA | UC | I.H. Lichter-Marck & J. Santore | August 2019 |
| *Laphamia lemmonii* A. Gray | ILM 458 | Big Hatchet Mountains, Hidalgo Co., NM, USA | UC | I.H. Lichter-Marck | August 2018 |
| *Laphamia vandevenderi* (B.H. Turner) I.H. Lichter-Marck | Vandevend er 2018- 87 | Sierra Cucurpe, Sonora, MX | US | T. Vandevende r & A.L. Reina | August 2017 |

| | | | | | |
|---|---|---|---|---|---|
| *Laphamia staurophylla* Barneby | ILM 802 | Truth or Consequences, NM., USA | UC | I.H. Lichter-Marck | September 2019 |
| *Laphamia staurophylla* Barneby | UC 1731838 | Fresnal Canyon, Alamagordo, Otero Co., NM, USA | UC | Dean Wm. Taylor | 21 July 1991 |
| *Laphamia rupestris* var. *rupestris* A. Gray | AZ 283476 | Davis Mountains, Davis Co., TX, USA | AZ | S. Sikes | September 1965 |
| *Laphamia fosteri* (A.M. Powell) I.H. Lichter-Marck | TEX 6528 | Panther Canyon, Apache Mountains, Culberson Co., TX, USA | TEX | J.F. Weedin | 1978 |
| *Laphamia fosteri* (A.M. Powell) I.H. Lichter-Marck | SRSU 1233 | Panther Canyon, Apache Mountains, Culberson Co., TX, USA | SRSU | A.M. Powell | 1978 |
| *Laphamia bisetosa* A. Gray | UC 1412751 | Maravillas Canyon, Brewster Co., TX, USA | UC | D.S. Correll & H.B. Correll | 9 November 1967 |
| *Laphamia cernua* Greene | ILM 393 | Organ Mountains, Doña Ana Co., NM, USA | UC | I.H. Lichter-Marck | September 2019 |
| *Laphamia lindheimeri* A. Gray | UC 1440574 | Loma Alta, Val Verde Co., TX, USA | UC | A.M. Powell & S. Sikes | 19 June 1965 |
| *Laphamia huecosensis* (A.M. Powell) I.H. Lichter-Marck | UC 1609992 | Sierra Juarez, Chihuahua, MX | UC | R. Spellenberg, L. Brouillet, D. Kearns. | 29 May 1993 |
| *Laphamia parryi* (A. Gray) Benth. & Hook. | ILM 377 | Chisos Mountains, Brewster Co., TX, USA | UC | I.H. Lichter-Marck & S. Winitsky | October 2018 |
| *Laphamia vaseyi* (J.M. Coult.) I.H. Lichter-Marck | AZ 416297 | Plateau south of Cigar Mountain, Brewster Co., TX, USA | AZ | Mark Fishbein; Angela Rein, Lindsey Worcester | 14 August 2013 |
| *Laphamia vaseyi* (J.M. Coult.) I.H. Lichter-Marck | ILM 355 | Ernst Tinaja, Brewster Co., TX, USA | UC | I.H. Lichter-Marck & S. Winitsky | October 2018 |

| | | | | | |
|---|---|---|---|---|---|
| *Laphamia dissecta* Torr. | Black s.n. | Lajitas, Presidio Co., TX, USA | SRSU | A. Black | 11 August 2019 |
| *Laphamia reinana* (B.L. Turner) I.H. Lichter-Marck | Santore s.n. | Sierra los Locos, Mncpio. Aconchi, Sonora, MX | UC | J. Santore | August 2019 |
| *Laphamia leptoglossa* (Harv. & A. Gray ex A. Gray) I.H. Lichter-Marck | MDE-2708 | Sierra la Campana, Hermosillo, Sonora, MX | SON | T.R. Vandevender | 25 December 2015 |
| *Laphamia cordifolia* (Rydb.) I.H. Lichter-Marck | AZ 347845 | Yecora, Sonora, MX | AZ | A.L. Reina-G.; T.R. Van Devender, T.F. Daniel, G.M. Ferguson, B.J. Syke | 17 March 1998 |
| *Laphamia cordifolia* (Rydb.) I.H. Lichter-Marck | UCR 80496 | Sierra de Alamos, Sonora, MX | UCR | A.C. Sanders | 20 March 1993 |
| *Galinsogeopsis coronopifolia* (A.Gray) I.H. Lichter-Marck | ILM 394 | Organ Mountains, Doña Ana Co., NM, USA | UC | I.H. Lichter-Marck & I. Nunez | October 2018 |
| *Galinsogeopsis pseudociliata* (A.M. Powell) I.H. Lichter-Marck | ARIZ356412 | Sierra La Brena on E slope of the Sierra Madre, Chihuahua, MX | AZ | J. Spencer; D. Atwood | 25 September 1998 |
| *Galinsogeopsis lineariloba* (Rydb.) I.H. Lichter-Marck | UC145620 | San Ramon, Durango, MX | UC | E. Palmer | 21 April-18 May 1906 |
| *Galinsogeopsis montana* (A.M. Powell) I.H. Lichter-Marck | UC 1523222 | Turuachi, Chihuahua, MX | UC | G. Nesom & P. Lewis | 24 Aug 1984 |
| *Galinsogeopsis montana* (A.M. Powell) I.H. Lichter-Marck | TEX 00141967 | Turuachi Canyon, Chihuahua, MX | TEX | G. Nesom & P. Lewis | 24 August 1984 |
| *Galinsogeopsis canescens* | UC 651747 | Sierra Tacuichamon | UC | H.S. Gentry | 12 February 1940 |

| | | | | | |
|---|---|---|---|---|---|
| (Everly) I.H. Lichter-Marck | | a, Sinaloa, MX | | | |
| *Galinsogeopsis spilanthoides* var. *spilanthoides* Sch. Bip. | CIAD 2652 | Isla Venados, Mazatlan, Sinaloa, MX | CIAD | M. Ruiz Guerrero. | 5 February 2017 |
| *Galinsogeopsis jaliscana* (A. Gray) I.H. Lichter-Marck | ILM 603 | Sierra de San Esteban, Jalisco, MX | UC | I.H. Lichter-Marck, P. Carrilo-Reyes, K. Soler, S. Winitsky | November 2018 |
| *Galinsogeopsis jaliscana* (A. Gray) I.H. Lichter-Marck | Lundell 60080 | El Diente de Sierra de San Esteban, Jalisco, MX | Lundell | A. García G., M. Dolores A. | 8 February 1994 |
| *Galinsogeopsis glaucescens* (B.H. Turner) I.H. Lichter-Marck | TEX 29908 | Juchipilas, Zacatecas, MX | TEX | J. Panero | 7 November 1996 |
| Galinsogeopsis feddemae (McVaugh) I.H. Lichter-Marck | NY 426075 | Valparaiso, Zacatecas, MX | NY | C. Feddema | 16 August 1969 |
| Galinsogeopsis sp. nov. "gamon" | | Sierra de Gamon, Durango, MX | CIIDIR | A. Castro-Castro | |
| *Nesothamnus incanus* (A. Gray) Rydb. | ILM | UCLA Botanical Garden, Los Angeles, CA, USA | UC | I.H. Lichter-Marck & E. Meyer | May 2019 |
| *Perityle rosei* Greenm. | ILM 621 | Sierra la Primavera, Jalisco, MX | UC | I.H. Lichter-Marck & P. Carrillo-Reyes | November 2018 |
| *Perityle rosei* Greenm. | CAS 694453 | Heujuquilla del Alto, Durango, MX | CAS | D.E. Breedlove, F. Almeda | 23 November 1983 |
| *Perityle trichodonta* S.F. Blake | ILM 615 | Bolaños, Jalisco, MX | UC | I.H. Lichter-Marck, P. Carrillo-Reyes, K. Soler, S. Winitsky | November 2018 |
| *Perityle trichodonta* S.F. Blake | UCR 70576 | Nayarit, a 22.7 km al SW de Jesus Maria, Camino a | UCR | O. Téllez V., G. Flores F. | 5 November 1988 |

| | | | | | |
|---|---|---|---|---|---|
| | | la Mesa del Nayar, MX | | | |
| *Pericome caudata* B.L. Rob. | ILM 118 | Silver Canyon, White Mountains, Inyo Co., CA, USA | UC | I.H. Lichter-Marck, K. Allerton, D. Neubauer | July 2018 |
| *Pericome caudata* B.L. Rob. | ILM 769 | Little Eden Spring, Humphrey's Peak, Coconino Co., AZ, USA | UC | I.H. Lichter-Marck | September 2018 |
| *Eutetras palmeri* A. Gray | UCR 117599 | Rincón de Ramos, Aguascalientes, MX | UCR | J.L. Villaseñor R. | 22 October 1995 |
| *Eutetras palmeri* A. Gray | UCR 117599 | Aguascalientes, MX | UCR | J.L. Villaseñor R. | October 22 1995 |
| *Perityle emoryi* Torr. | UC 648836 | Caldera, Atacama Province, Chile | UC | A.A. Beetle | 19 February 1939 |
| *Perityle emoryi* Torr. | ILM2017-3 | Punta San Jose, Baja California, MX | UC | I.H. Lichter-Marck | May 2017 |
| *Perityle emoryi* Torr. | ILM 693 | Cataviña, Baja California, MX | UC | I.H. Lichter-Marck | January 2019 |
| *Perityle emoryi* Torr. "yellow" | JEPS 128783 | Chocolate Mountains, Imperial Co., CA, USA | UC | A. Sanders, J. Malusa, & I.H. Lichter-Marck | March 2017 |
| *Perityle emoryi* Torr. | ILM 191 | Hanuapah Canyon, Inyo Co., CA, USA | UC | I.H. Lichter-Marck & K. Allerton | June 2018 |
| *Perityle socorrensis* Rose | SD 259737 | Isla San Benedicto, Colima, MX | SD | J. Rebman, E. Excurra, S.Vanderplank | |
| *Perityle socorrensis* Rose | SD 259739 | Isla Clarion, Colima, MX | SD | J. Rebman, E. Excurra, S. Vanderplank | |
| *Lycapsus tenuifolius* Phil. | SD | Isla San Felix, Desventuradas Islands, Chile | SD | R. Thorne | |

| | | | | | |
|---|---|---|---|---|---|
| *Amauria brandegeeana* (Rose) Rydb. | UCR 146783 | Baja California Sur, MX | UCR | N/A | |
| *Perityle californica* Benth. | Reina 2019-04 | Cerro Colorado, Sonora, MX | US | A.L. Reina & T. Vandevender | Jan 2019 |
| *Perityle californica* Benth. | ILM 692 | San Ignacio, Baja California, MX | UC | I.H. Lichter-Marck | Jan 2019 |
| *Perityle californica* Benth. | UC 1787616 | Ciudad Constitucion, Baja California Sur, MX | UC | J.P. Rebman | 24 October 2001 |
| *Perityle cuneata* var. *marginata* (Rydb.) I.M. Johnst. | UC 1116153 | Playa el Coyote, La Paz, Baja California Sur, MX | UC | D.M. Porter | 30 December 1958 |

**SI References**

Sample References:

1. J.J. Doyle, J.L. Doyle, CTAB DNA extraction in plants. *Phytochemical Bulletin,* 19, 11–15 (1987).
2. I.H. Lichter-Marck, W.A. Freyman, C.M. Siniscalchi, J.R. Mandel, A. Castro-Castro, G. Johnson, B.G. Baldwin, Phylogenomics of Perityleae (Compositae) provides new insights into morphological and chromosomal evolution of the rock daisies. *J. Syst. Evol.*, 58(6), 853-880 (2020).
3. J.R. Mandel, R.B. Dikow, V.A. Funk, R.R. Masalia, S.E. Staton, A. Kozik, R.W. Michelmore, L.H. Rieseberg, J.M. Burke, A target enrichment method for gathering phylogenetic information from hundreds of loci: An example from the Compositae. *Applications in Plant Sciences*, 2(2), 1300085 (2014).
4. J.R. Mandel, M.S. Barker, R.J. Bayer, R.B. Dikow, T.G. Gao, K.E. Jones, S. Keeley, N. Kilian, H. Ma, C.M. Siniscalchi, A. Susanna, The Compositae tree of life in the age of phylogenomics. *J. Syst. Evol.*, 55(4), 405-410 (2017).
5. J.R. Mandel, R.B. Dikow, V.A. Funk, Using phylogenomics to resolve mega-families: An example from Compositae. *J. Syst. Evol.*, 53(5), 391-402 (2015).
6. A.M. Kozlov, D. Darriba, T. Flouri, B. Morel, A. Stamatakis, RAxML-NG: A fast, scalable and user-friendly tool for maximum likelihood phylogenetic inference. *Bioinformatics*, 35(21), 4453-4455 (2019).
7. A.J. Aberer, K. Kobert, A. Stamatakis, ExaBayes: Massively parallel Bayesian tree inference for the whole-genome era, *Mol. Biol. Evol.*, 31(10), 2553-2556 (2014).
8. C. Zhang, E. Sayyari, S. Mirarab, “ASTRAL-III: increased scalability and impacts of contracting low support branches.” In *RECOMB International Workshop on Comparative Genomics*, pp. 53-75, (Springer, Cham., 2017).
9. M. Dos Reis, P.C. Donoghue, Z. Yang, Bayesian molecular clock dating of species divergences in the genomics era. *Nat. Rev. Gen.*, 17(2), 71-80 (2017).
10. M. Dos Reis, J. Inoue, M. Hasegawa, R.J. Asher, P.C. Donoghue, Z. Yang, Phylogenomic datasets provide both precision and accuracy in estimating the timescale of placental mammal phylogeny. *Proc. Biol. Sci.*, 279(1742), 3491-3500 (2012).
11. A. Rambaut, A.J. Drummond, D. Xie, G. Baele, M.A. Suchard, Posterior summarization in Bayesian phylogenetics using Tracer 1.7. *Syst. Biol.*, 67(5), 901 (2018).
12. N.M. Koch, Phylogenetic subsampling and the search for phylogenetically reliable loci. *Mol. Biol. Evol.,* 38(9), 4025-4038 (2021).
13. S. Höhna, M.J. Landis, T.A. Heath, B. Boussau, N. Lartillot, B.R. Moore, J.P. Huelsenbeck, F. Ronquist, RevBayes: Bayesian phylogenetic inference using graphical models and an interactive model-specification language. *Syst. Biol.*, 65(4), 726-736 (2016).
14. E.E. Goldberg, B. Igić, On phylogenetic tests of irreversible evolution. *Evolution*, 62(11), 2727-2741 (2008).
15. B.G. Baldwin, B.L. Wessa, J.L. Panero, Nuclear rDNA evidence for major lineages of helenioid Heliantheae (Compositae). *Syst. Bot.*, 161-198 (2002).
16. M. Pagel, A. Meade, Bayesian analysis of correlated evolution of discrete characters by reversible-jump Markov chain Monte Carlo. *Am. Nat.*, *167*(6), 808-825 (2006).
17. M. Pagel, Detecting correlated evolution on phylogenies: A general method for the comparative analysis of discrete characters. *Proc. Biol. Sci.,* 255(1342), 37-45 (1994).
18. W. Xie, P.O. Lewis, Y. Fan, L. Kuo, M.H. Chen, Improving marginal likelihood estimation for Bayesian phylogenetic model selection. *Syst. Biol.*, 60(2), pp.150-160 (2011).
19. J. Rzedowski, *The vegetation of Mexico*. (Editorial Limusa, 1981).
20. J. McPhee, *Annals of the former world*. (Farrar, Straus and Giroux, 2000).
21. I.L. Wiggins, *Flora of Baja California*. (Stanford University Press, 1980).
22. J. P. Rebman, J. Gibson, K. Rich. Annotated checklist of the vascular plants of Baja California, Mexico. (San Diego Natural History Society, 2016).
